# Break-induced replication is activated to repair R-loop-associated double-strand breaks in SETX-deficient cells

**DOI:** 10.1101/2024.06.29.601361

**Authors:** Tong Wu, Youhang Li, Linda Z. Shi, Xiaohua Wu

## Abstract

The primary role of break-induced replication (BIR) is to repair single-ended double-strand breaks (seDSBs) generated at broken replication forks and eroding telomeres. In this study, we demonstrated that when senataxin (SETX), an RNA/DNA helicase, is defective, hyper-recombination using the BIR mechanism is induced at R-loops/hybrids-accumulated double-ended DSBs (deDSBs), uncovering a role for BIR in repair of R-loops/hybrids-associated deDSBs. Intriguingly, the loss of SETX not only triggers non-canonical hyper-end resection requiring RAD52 and XPF, but also stalls Polα-primase-initiated end-fill DNA synthesis due to the accumulation of RNA/DNA hybrids on single-strand DNA (ssDNA) overhangs at deDSBs. This conflict between fill-in DNA synthesis and accumulated hybrids induces PCNA ubiquitination and PIF1 loading, thereby initiating the BIR mechanism at deDSBs. Hyper-resection further enhances PCNA ubiquitination and PIF1 loading, driving BIR-mediated hyper-recombination. Moreover, SETX is synthetic lethal with PIF1, RAD52, and XPF, offering new strategies for targeted treatment of SETX-deficient tumors.

## Introduction

DNA double-strand breaks (DSBs) represent the most dangerous form of DNA damage, which could lead to genome instability and, more severely, to cell death (Khanna and Jackson, 2001). DSBs can be repaired by two major pathways; homologous recombination (HR) and non-homologous end joining (NHEJ) (Ceccaldi et al., 2016). Homology-dependent HR serves as an error-free mechanism to repair DSBs (Heyer, 2015; Jasin and Rothstein, 2013; Paques and Haber, 1999), whereas homology-independent NHEJ could be error-prone in some cases (Zhao et al., 2020). While NHEJ remains active throughout the cell cycle, HR is activated when cells enter S and G2 phases when sister chromatids are available to serve as templates (Hustedt and Durocher, 2016). Break-induced replication (BIR) is a subtype of HR, which operates at one-ended DSBs (seDSBs), such as collapsed replication forks and eroding telomeres (Anand et al., 2013; Liu and Malkova, 2022; Llorente et al., 2008; Wu and Malkova, 2021a). BIR and HR share similar initiation steps, including end resection and strand invasion, but diverge at the DNA synthesis step, where BIR exhibits a much longer DNA synthesis tract than HR (Kramara et al., 2018). To achieve long-tract DNA synthesis, BIR employs a unique feature distinct from HR; BIR replication is dependent on POLD3/Pol32, a subunit of DNA polymerase δ, and helicase PIF1 (Donnianni and Symington, 2013; Li et al., 2021a; Lydeard et al., 2007; Saini et al., 2013; Wilson et al., 2013). Notably, BIR is mutagenic, which is often associated with template switching (Sakofsky et al., 2015; Smith et al., 2007) and exhibits hyper mutation rate (Sakofsky et al., 2012; Sakofsky et al., 2014). Thus, BIR needs to be tightly regulated, and its activation should only occur when its use is necessary. In yeast, BIR has been shown to be promoted under conditions such as defects in second end capture, persistent DNA synthesis within D-loops, unbalanced end resection of the two DSB ends, and impaired DNA damage checkpoint (Pham et al., 2021). In mammalian cells, 53BP1 has been found to suppress BIR (Shah et al., 2024); however, the regulation of BIR activation through this pathway under physiological and pathological conditions remains unclear.

R-loops are three-stranded nucleic acid structures formed by extended annealing of nascent RNA to the DNA template, composing an RNA/DNA hybrid and displaced single-strand DNA (ssDNA). While R-loops possess important biological functions such as transcription regulation, DNA repair, and maintaining telomere integrity, their persistence can be detrimental, leading to genome instability (Garcia-Muse and Aguilera, 2019; Niehrs and Luke, 2020). Abundant evidence has suggested that R-loops and RNA/DNA hybrids (R-loops/hybrids) are induced at DSB sites and actively involved in DSB repair (Brickner et al., 2022; Gomez-Gonzalez and Aguilera, 2023; Marnef and Legube, 2021; Petermann et al., 2022), but a comprehensive understanding of the underlying mechanisms is still elusive. Some studies suggest that R-loops/hybrids formed at DSB ends interfere with the repair process, while others show that R-loops/hybrids have a stimulatory effect on repair. Loss of R-loop resolution factors such as DDX5, DHX9, and HNRNPD causes R-loop accumulation at DSBs, impeding end resection and causing defective HR (Alfano et al., 2019; Matsui et al., 2020; Sessa et al., 2021; Yu et al., 2020), whereas increased R-loop formation due to inactivation of the exosome catalytic subunit EXOSC10 strongly stimulates end resection (Domingo-Prim et al., 2019). Moreover, DSB-induced small RNAs (diRNAs) and damage-induced long non-coding RNAs (dilncRNAs) facilitate the recruitment of HR proteins to DSBs (D’Alessandro et al., 2018; Gao et al., 2014; Michelini et al., 2017). In G2 cells exposed to IR, RAD52 is recruited to DNA damaging sites in a transcription- and R-loop-dependent manner to promote HR (Yasuhara et al., 2018). In yeast, defects in both forming and removing RNA/DNA hybrids reduce HR efficiency, suggesting that transient RNA/DNA hybrid formation at DSBs is essential for HR (Ohle et al., 2016). These findings suggest that different repair outcomes may result from the complex and intricate regulation exerted by R-loops/hybrids at different DSB repair steps, involving multiple players. Understanding the detailed mechanisms by which R-loop-resolving factors modulate DSB repair is therefore important for uncovering the biological significance of R-loops/hybrids at DSB ends.

SETX encodes a putative RNA-DNA helicase capable of unwinding RNA-DNA hybrids and plays an important role in R-loop homeostasis (Gatti et al., 2023; Giannini and Porrua, 2024; Groh et al., 2017). Human SETX defects are documented in the neurodegenerative disorder ataxia with oculomotor apraxia 2 (AOA-2) (Moreira et al., 2004) and autosomal dominant amyotrophic lateral sclerosis type 4 (ALS4) (Chen et al., 2004). Substantial evidence suggests an important role of SETX in the maintenance of genome stability (Gatti et al., 2023; Giannini and Porrua, 2024; Groh et al., 2017). SETX and its yeast orthologue Sen1 interact with RNA Pol II (Chen et al., 2006; Suraweera et al., 2009; Ursic et al., 2004) and are involved in transcription regulation and termination (Gatti et al., 2023; Giannini and Porrua, 2024). SETX and Sen1 associate with replication forks, promoting replication through transcription-active genes (Alzu et al., 2012). SETX forms foci colocalizing with DNA repair proteins upon replication stress and is believed to participate in resolving transcription and replication conflict (TRC) (Richard et al., 2013; Yuce and West, 2013). SETX in complex with BRCA1 is also involved in dismantling R-loops at transcription pause sites in transcription termination regions to prevent DNA damage (Hatchi et al., 2015). Furthermore, it has been demonstrated that SETX and Sen1 are recruited to DSBs when they occur in transcriptionally active loci to limit accumulation of R-loops/hybrids at DSB ends (Cohen et al., 2022; Cohen et al., 2018; Hatchi et al., 2015; Rawal et al., 2020).

In this study, using EGFP-based repair reporters, which specifically monitor HR and BIR pathways in mammalian cells (Li et al., 2021b), we detected hyper-recombination activity in SETX-deficient cells. Interestingly, short tract gene conversion (STGC), which is usually referred to as HR, alters its genetic dependence in SETX-deficient cells and shows characteristics of BIR, requiring PIF1 and POLD3. End resection is substantially stimulated in an R-loop-dependent manner at DSBs using an unconventional mechanism when SETX is deficient. Accumulated RNA/DNA hybrids at DSB ends cause stalling of end-fill DNA synthesis on ssDNA overhangs, resulting in PCNA ubiquitination and PIF1 recruitment, which activates the BIR mechanism. The shift in the repair mechanism from HR to BIR at deDSBs in SETX-deficient cells makes them reliant on the BIR pathway for survival, providing potential therapeutic strategies for targeting SETX-deficient tumors.

## Results

### SETX loss induces hyper-recombination for both HR and BIR in an R-loop-dependent manner

In our previous study, we established EGFP-based reporter systems to determine the efficiency of DSB repair mediated by the HR (STGC) pathway (Figure 1A) and the BIR (long tract gene conversion: LTGC) pathway (Figure 1B) (Li et al., 2021b). In yeast, gene conversion with tract length longer than 1 kb, uses the BIR mechanism (Jain et al., 2009; Mehta et al., 2017). In our EGFP-BIR reporter, green cells are produced only when DNA synthesis extends 1.2 kb to 3.8 kb beyond the FP region, which is terminated by either end joining (BIR-EJ) or by synthesis-dependent strand annealing (SDSA, BIR-SDSA) (Figure S1A) (Li et al., 2021b). Interestingly, we found that SETX depletion or knockout (KO) results in significantly higher percentages of EGFP-positive cells in both U2OS (EGFP-HR) and U2OS (EGFP-BIR) reporter cell lines after I-SceI cleavage (Figure 1C and 1D), suggesting that SETX deficiency induces hyper-recombination for both HR and BIR. To determine whether the helicase activity of SETX is required for suppressing hyper-recombination, we expressed SETX wildtype (WT) allele or helicase-dead mutant SETX-P-loopΔ, containing deletion of eight amino acids (Δ1963-1970aa) in the SETX P-Loop GTP/ATP binding motif (Bennett et al., 2020) in both WT and *SETX*-KO U2OS (EGFP-BIR) reporter cells. While expressing the SETX-WT allele suppresses hyper-BIR in *SETX*-KO cells, the helicase-dead mutant failed to do so (Figure 1E). Consistent with the role of SETX in resolving R-loops (Crossley et al., 2023; Zhao et al., 2022), we showed stronger R-loop staining in *SETX*-KO cells compared to WT cells, with RNASEH1 overexpression attenuating the increase (Figure S1D). Overexpression of RNASEH1 in SETX-depleted cells also leads to suppression of hyper-recombination for BIR (Figure 1F) and HR (Figure 1G). These results suggest that SETX deficiency promotes hyper-recombination for both HR and BIR in an R-loop-dependent manner.

**Figure 1.**
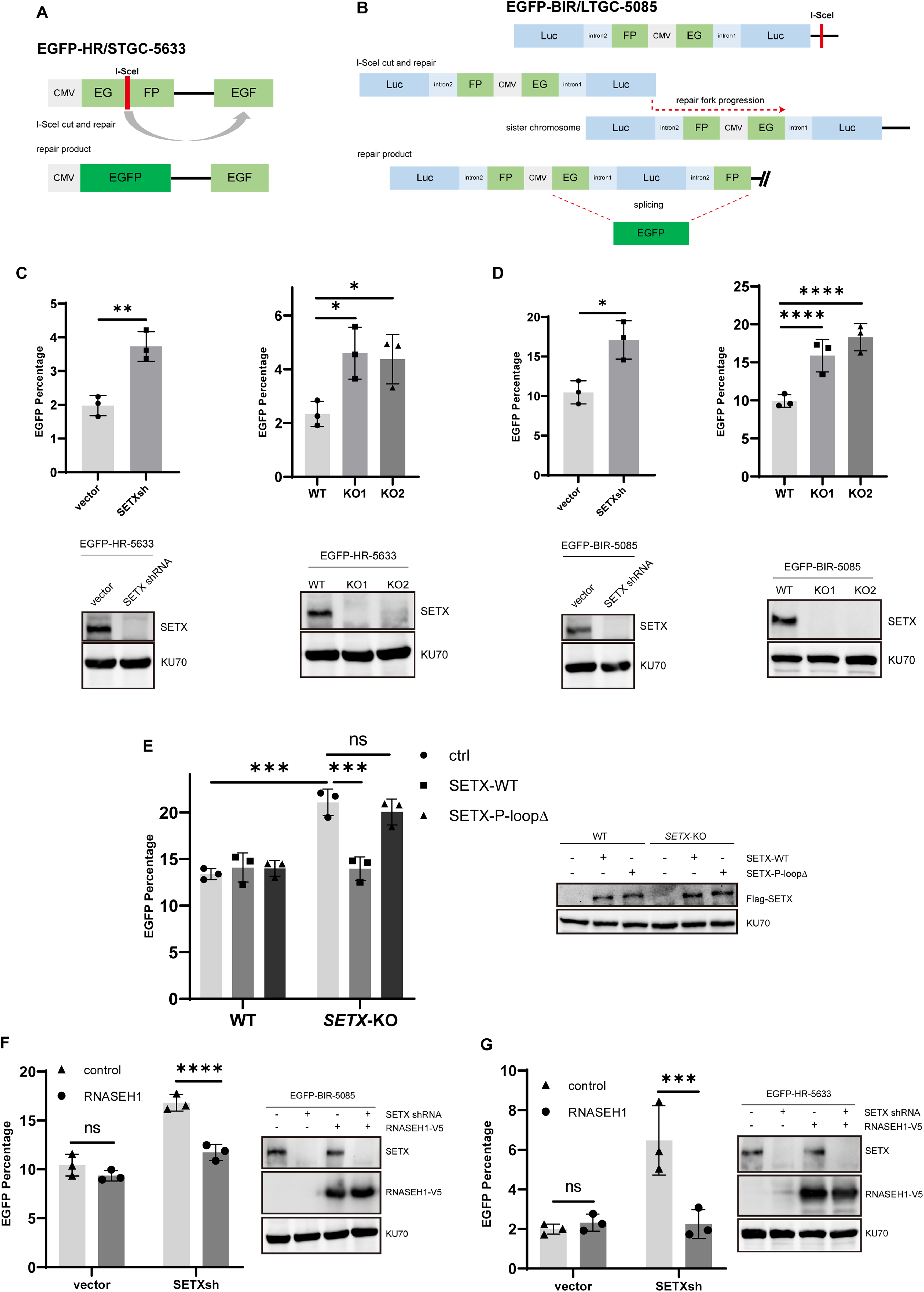
SETX loss induces hyper-recombination for both HR and BIR in an R-loop-dependent manner. A, B. Schematic drawing of EGFP-HR-5633 (A) and EGFP-BIR-5085 (B) reporters. C, D. U2OS WT and *SETX*-KO EGFP-HR-5633 (C) and EGFP-BIR-5085 (D) reporter cells expressing SETX shRNA or vector, were infected with lentivirus expressing I-SceI to induce DSBs. The percentage of EGFP-positive cells was quantified by FACS analysis five days later (top). The efficiency of SETX knockdown or knockout were determined by Western blotting (bottom). E. U2OS WT or *SETX*-KO EGFP-BIR-5085 reporter cells were infected with lentivirus expressing Flag-tagged WT SETX or mutated SETX, then infected with lentivirus expressing I-SceI to induce DSBs. The percentage of EGFP-positive cells was quantified by FACS analysis five days later (Left). The expression of Flag-SETX was determined by Western blotting (Right). F, G. U2OS EGFP-HR-5633 (G) or EGFP-BIR-5085 (F) reporter cells with or without RNASEH1 expression were infected with lentiviruses expressing SETX shRNA or vector. Three days later, all cells were infected with lentiviruses expressing I-SceI to induce DSBs. The percentage of EGFP-positive cells was quantified by FACS analysis five days later (Left). The expression level of SETX and RNASEH1-V5 was determined by Western blotting (Right).

### Hyper-recombination induced in SETX-deficient cells is mediated by the BIR mechanism

It has been well established that PIF1 helicase is required for BIR (LTGC) but is dispensable for HR (STGC) (Li et al., 2021b). We showed that BIR in WT cells and hyper-BIR in *SETX*-KO cells are both strongly impaired upon PIF1 depletion by shRNAs, confirming the use of BIR (Figure 2A and Figure S2A). Interestingly, however, while HR (STGC) in SETX-proficient WT cells is not dependent on PIF1, depleting PIF1 in *SETX*-KO cells leads to a significant reduction in HR, suggesting that in SETX-deficient cells, PIF1 becomes indispensable for HR (STGC) (Figure 2B and S2B). This suggests that SETX deficiency disrupts the regulation of HR and BIR onset and induces hyper-recombination at deDSBs, using the BIR-like mechanism that requires PIF1.

**Figure 2.**
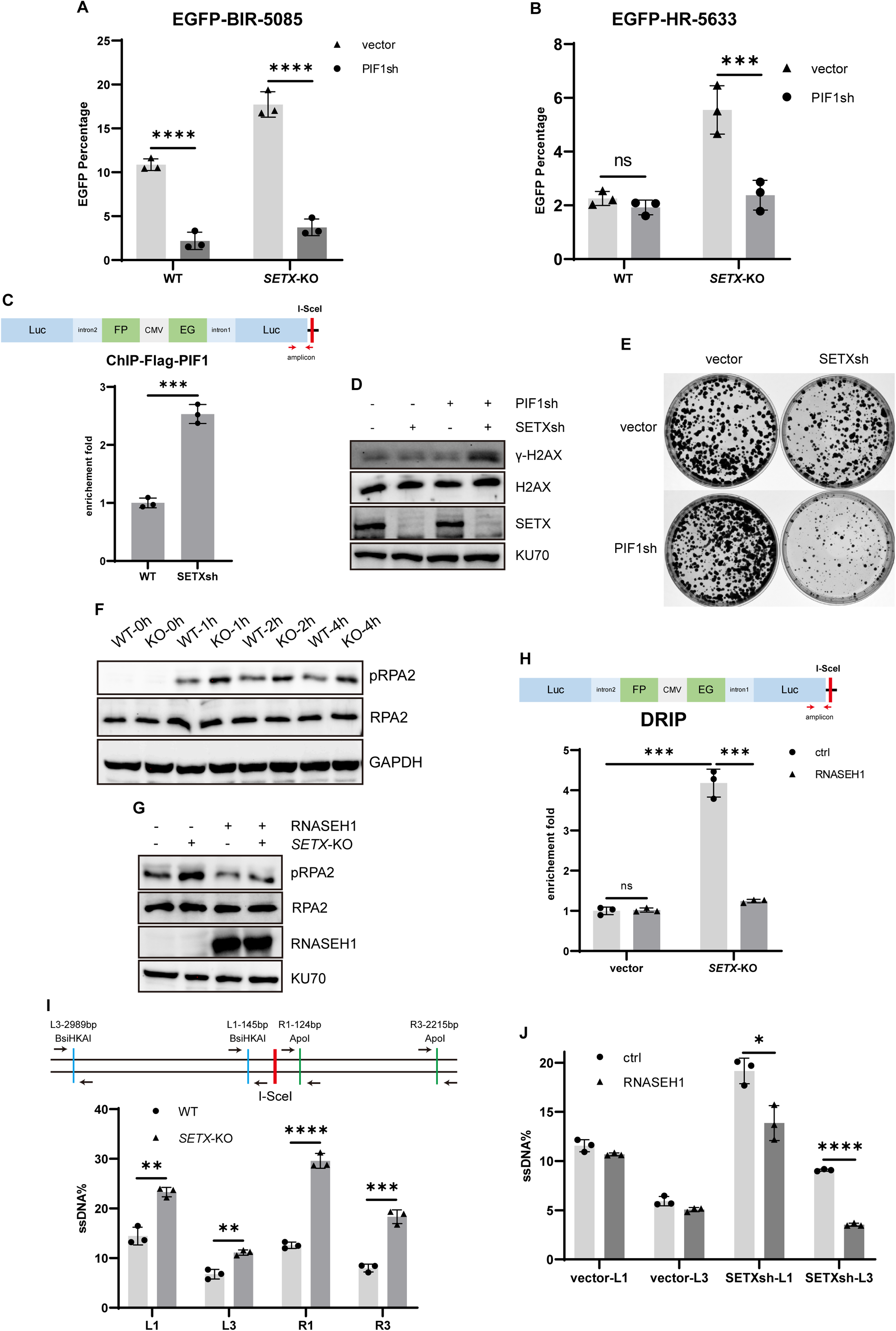
HR in SETX-deficient cells utilizes BIR-like mechanism. A, B. U2OS WT and *SETX*-KO EGFP-BIR-5085 reporter cells (A) or EGFP-HR-5633 reporter cells (B) were infected with lentiviruses expressing PIF1 shRNA or vector. After three days, these cells were infected with I-SceI-expressing lentiviruses to induce DSBs. The percentage of EGFP-positive cells was quantified by FACS analysis five days later. C. U2OS WT EGFP-BIR-5085 reporter cells were infected with Flag-PIF1 lentiviruses. Cells with Flag-PIF1 expression were infected with SETX shRNA or vector. Three days later, cells were further infected with I-SceI-expressing lentivirus to induce DSBs. Two days after infection, cells were harvested for ChIP analysis of Flag-PIF1 at indicated locus. D. U2OS cells were infected with lentiviruses expressing vector, SETX shRNA, PIF1 shRNA, or both SETX and PIF1 shRNA. Cell lysates were collected four days after lentiviral infection and analyzed by Western blotting for γH2AX and SETX, using KU70 as a loading control. E. U2OS cells were infected with lentiviruses expressing vector, SETX shRNA, PIF1 shRNA, or both SETX and PIF1 shRNAs. Three days after infection, cells were resuspended, counted, and 5000 cells were seeded on plates. After two weeks of culture, cells were stained with crystal violet. F. WT and *SETX*-KO U2OS cells were treated with 10 Gy IR. Cell lysates were collected at indicated time points and Western blotting was performed using indicated antibodies. G. WT and *SETX*-KO U2OS cells with or without RNASEH1 expression were treated with 10 Gy IR. Cell lysates were collected one hour after IR and Western blotting was performed using indicated antibodies. H. U2OS WT and *SETX*-KO EGFP-BIR-5085 reporter cells, with or without RNASEH1 expression, were infected with I-SceI-expressing lentiviruses to induce DSBs. Two days after infection, cells were harvested for DRIP analysis at indicated locus. I. Illustration of site-specific end resection assay in EGFP-BIR-5085 reporter (top). U2OS WT and *SETX*-KO EGFP-BIR-5085 reporter cells were infected with I-SceI-expressing lentiviruses to induce DSBs (bottom). After two days, genomic DNA (gDNA) is extracted and digested with indicated restriction enzyme (RE): BsiHKAI for the left DSB ends, ApoI for the right DSB ends, or SacII as mock for both ends. RE-resistant ssDNA is quantified by qPCR using primer sets around the RE recognition sites. J. U2OS EGFP-BIR-5085 reporter cells with or without RNASEH1 expression were infected with SETX shRNA or vector. Three days after infection, cells were infected with I-SceI-expressing lentivirus to induce DSBs. Two days after the second infection, cells were harvested for gDNA for end resection assay.

BIR is responsible for repairing seDSBs on broken forks (Kramara et al., 2018; Wu and Malkova, 2021a). We previously proposed that the assembly of BIR replisomes, containing PIF1 and requiring POLD3 activity, triggers the onset of the BIR mode, and once assembled, BIR replisomes can facilitate both HR/STGC and BIR/LTGC (Li et al., 2021a). At broken forks, BIR replisomes are readily assembled and facilitate both BIR/LTGC and HR/STGC to repair seDSBs, although BIR/LTGC is utilized more frequently (Figure S1B left) (Li et al., 2021a). However, at deDSBs, such as those generated by endonucleases and ionizing radiation (IR), BIR replisomes are assembled only when BIR is in need, and thus PIF1- and POLD3-independent HR/STGC is utilized predominantly over PIF1- and POLD3-dependent BIR/LTGC (Li et al., 2021a) (Figure S1B right). We propose that loss of SETX triggers the assembly of BIR replisomes at deDSBs, thereby activating the BIR mode even for HR/STGC (Figure S1C), reminiscent of BIR activation at seDSBs on broken forks for both HR/STGC and BIR.

We performed chromatin immunoprecipitation (ChIP) at DSBs generated by I-SceI cleavage in the BIR reporter, and observed that SETX depletion causes increased recruitment of Flag-PIF1 to DSBs generated by I-SceI cleavage in the BIR reporter (Figure 2C). We speculate that enhanced PIF1 recruitment to DSBs in SETX-deficient cells may facilitate the onset of the BIR mechanism for DSB repair. Additionally, we observed that γ-H2AX is significantly increased in cells depleted for both PIF1 and SETX (Figure 2D), and SETX and PIF1 exhibit synthetic lethal interactions (Figure 2E). Collectively, these data suggest that hyper-recombination induced in SETX-deficient cells primarily occurs through activating the BIR mechanism rather than HR, and thus BIR becomes an essential DSB repair mechanism for the survival of SETX-deficient cells.

### SETX loss induces excessive end resection mediated by a non-canonical pathway, contributing to hyper-recombination

The impact of accumulated R-loops on DSB repair often involves the regulation of end resection (Alfano et al., 2019; Domingo-Prim et al., 2019; Matsui et al., 2020; Sessa et al., 2021; Yu et al., 2020). To investigate the effect of SETX loss on end resection, we first examined the level of RPA2 phosphorylation at S4 and S8 (pRPA2) after ionizing radiation (IR), known to be induced by end resection (Liaw et al., 2011). *SETX*-KO cells manifest stronger pRPA2 signals after IR compared to WT cells (Figure 2F), indicative of overactive end resection. Overexpression of RNASEH1 suppresses increased pRPA2 signals in *SETX*-KO cells, suggesting that hyper-resection is indeed caused by accumulation of R-loops/hybrids when SETX is deficient (Figure 2G).

By DNA/RNA immunoprecipitation (DRIP) analysis, we showed that R-loops/hybrids are accumulated at the DSBs in our BIR reporter, which can be suppressed by overexpressing RNASEH1 (Figure 2H). By using the well-established quantitative end resection assay (Zhou et al., 2014), we demonstrated that end resection is significantly enhanced in *SETX*-KO cells at I-SceI-induced DSBs in the BIR reporter (Figure 2I), and overexpression of RNASEH1 suppresses hyper-resection due to SETX deficiency (Figure 2J). Pretreating genomic DNA with RNase H prior to end resection assay leads to a similar observation that loss of SETX results in hyper-resection (Figure S2C), excluding the possibility that forming R-loops/hybrids at DSBs hinders end resection assay. These data suggest that hyper-resection in SETX-deficient cells is induced by accumulated R-loops/hybrids at DSBs.

To investigate how hyper-resection is induced in SETX-deficient cells, we examined the role of MRE11 and CtIP in promoting hyper-resection. Surprisingly, after I-SceI cleavage, hyper-resection observed in *SETX*-KO cells is largely dependent on MRE11 but not CtIP (Figure 3A). Similarly, IR-induced pRPA2 is dependent on MRE11 but not CtIP in *SETX*-KO cells, while RPA2 phosphorylation requires both MRE11 and CtIP in WT cells (Figure 3B). It is well established that the MRE11-RAD50-NBS1 (MRN) complex initiates the end resection by using the endonuclease activity of MRE11 to make a nick on the 5’ strand with some distance from the break end, followed by using the 3’ to 5’ exonuclease activity of MRE11 to degrade DNA from the nick towards the DSB end (Cejka and Symington, 2021; Deshpande et al., 2016; Shibata et al., 2014). It has also been shown that CtIP is important for stimulating the MRE11 endonuclease activity (Anand et al., 2019; Anand et al., 2016; Cannavo and Cejka, 2014; Zdravkovic et al., 2021). The dispensable role of CtIP for hyper-resection in SETX-deficient cells implies that the initial MRN-mediated endonuclease cleavage at DSB ends may not be involved. To test this possibility, we used an inhibitor PFM01 that specifically inhibits the endonuclease activity of MRE11 (Shibata et al., 2014). Significantly, treatment with PFM01 results in a reduction of end resection in SETX-proficient cells but fails to do so in SETX-deficient cells, and instead even stimulates it, while XPF depletion still leads to reduction of end resection in the presence of PFM01 in *SETX*-KO cells but not in WT cells (Figure 3C). On the other hand, when we used the inhibitor PFM39, which specifically inhibits the exonuclease activity of MRE11 (Shibata et al., 2014), we observed a clear reduction of end resection close to the cleavage site in both WT and *SETX*-KO cells (Figure 3D left, L1, 145 bp left to I-SceI site, Figure 3D top). However, at the site L3, which is further away from the cleavage site (L3, 2989 bp left to I-SceI site, Figure 3D top), PFM39 treatment causes a minor decrease of end resection in *SETX*-KO cells with almost no effect in WT cells (Figure 3D right), consistent with the notion that extended 5’ to 3’ end resection involves BLM/DNA2 and EXO1, but not the exonuclease activity of MRE11. Collectively, these results suggest that the initiation of abnormal hyper-resection in SETX-deficient cells is not through the canonical pathway that requires MRE11- and CtIP-dependent endonucleolytic cleavage at DSB ends. However, the exonuclease activity of MRE11 is still required for 3’ to 5’ strand removal towards the DSB ends (Figure 7A).

**Figure 3.**
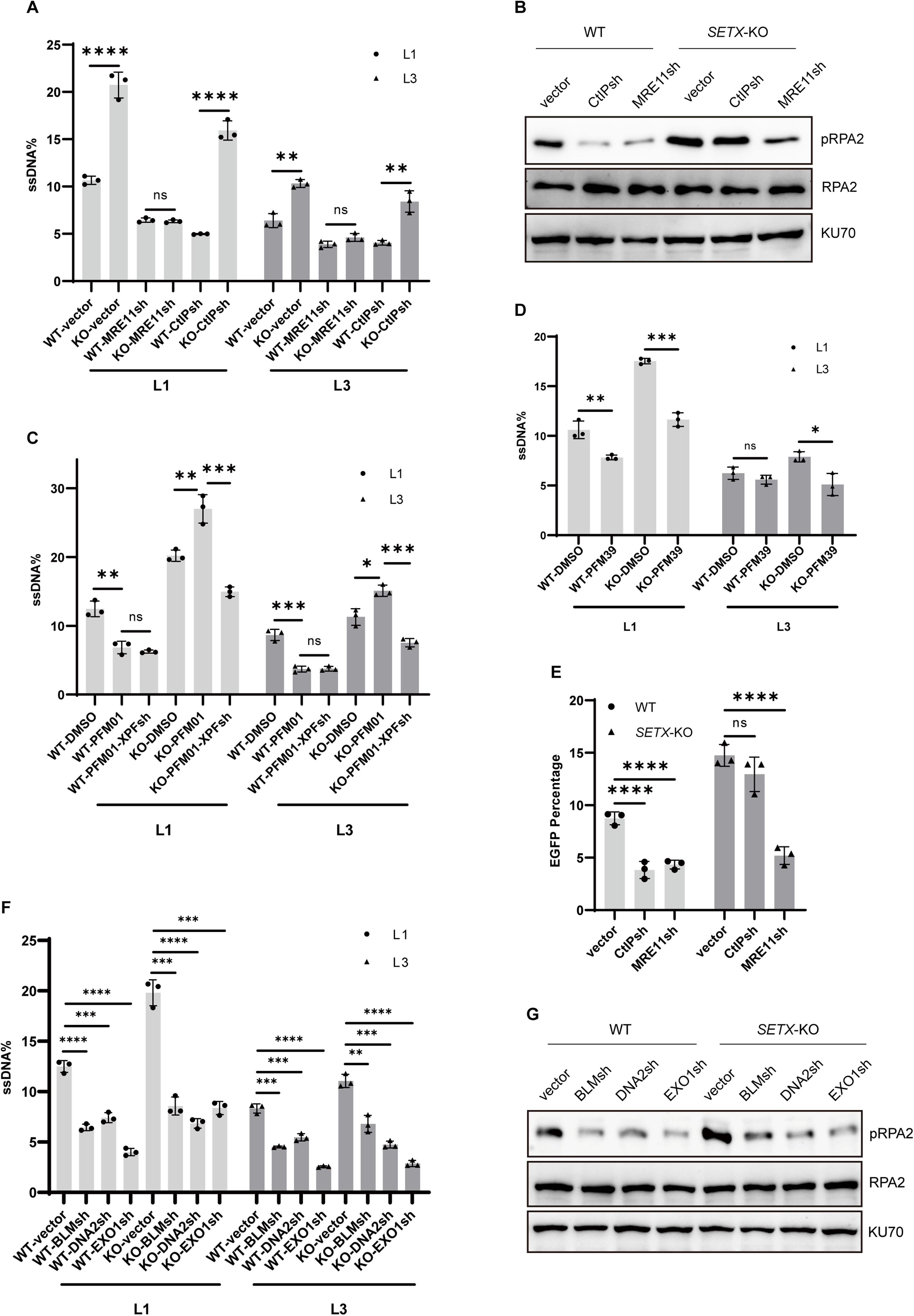
SETX loss induces hyper-resection through a non-canonical resection pathway. A, B, E. U2OS WT and *SETX*-KO EGFP-BIR-5085 reporter cells were infected with lentiviruses expressing MRE11 shRNA, CtIP shRNA, or vector. Three days were given to allow shRNA expression to achieve effective knockdown of each gene. For end resection assay (A), shRNA-treated cells were infected with I-SceI-expressing lentiviruses to induce DSBs. Two days after I-SceI infection, cells were harvested for end resection assay. For examining the pRPA2 level (B), shRNA-treated cells were treated with 10 Gy IR. Cell lysates were collected one hour after IR and Western blotting was performed using indicated antibodies. For BIR efficiency analysis (E), shRNA-treated cells were infected with I-SceI-expressing lentiviruses to induce DSBs. The percentage of EGFP-positive cells was quantified by FACS analysis five days after I-SceI infection. C. U2OS WT and *SETX*-KO EGFP-BIR-5085 reporter cells were infected with lentiviruses expressing XPF shRNA or vector. Three days later, cells were infected with I-SceI-expressing lentiviruses to induce DSBs. Inhibitor PFM01 (100 uM) or DMSO was applied to cells together with I-SceI lentiviruses. Two days after I-SceI infection, cells were harvested for end resection assay. D. U2OS WT and *SETX*-KO EGFP-BIR-5085 reporter cells were infected with I-SceI-expressing lentiviruses to induce DSBs, simultaneously treated with DMSO or inhibitor PFM39 (100 uM). Two days after I-SceI induction, cells were harvested for end resection assay. F, G. U2OS WT and *SETX*-KO EGFP-BIR-5085 reporter cells were infected with lentiviruses encoding BLM shRNA, DNA2 shRNA, EXO1 shRNA, or vector. Three days later, cells were then infected with I-SceI-expressing lentiviruses to induce DSBs for end resection assay (F), or treated with 10 Gy IR to examine the pRPA2 level (G).

We also performed BIR assay after depleting MRE11 or CtIP. While BIR is impaired in SETX-proficient cells after silencing CtIP or MRE11, BIR in *SETX*-KO cells is disrupted only after depleting MRE11 but not CtIP (Figure 3E, S3A and S3B). As CtIP is dispensable but MRE11 is required for hyper-resection in SETX-deficient cells, these findings reveal a correlation between hyper-resection and hyper-BIR, suggesting that the hyper-resection observed in the absence of SETX likely contributes to the promotion of hyper-BIR.

### R-loop accumulation at DSBs triggers RAD52- and XPF-dependent hyper-resection in SETX-deficient cells

The observation that in SETX-deficient cells, CtIP and the endonuclease activity of MRE11 are dispensable for end resection suggests the involvement of a non-canonical end resection mechanism. Interestingly, depleting flap endonuclease XPF leads to a reduction of BIR in *SETX*-KO cells but not in WT cells (Figure 4A and S4A). The end resection assay further demonstrated that hyper-resection in *SETX*-KO cells, but not normal end resection in SETX-proficient WT cells, is dependent on XPF (Figure 4B). XPF has been shown to physically interact with RAD52 (Motycka et al., 2004), and RAD52 is recruited to DSBs containing R-loops at transcriptionally active loci (Tan et al., 2020; Yasuhara et al., 2018). We speculate that XPF may be recruited through the R-loop-RAD52 axis.

**Figure 4.**
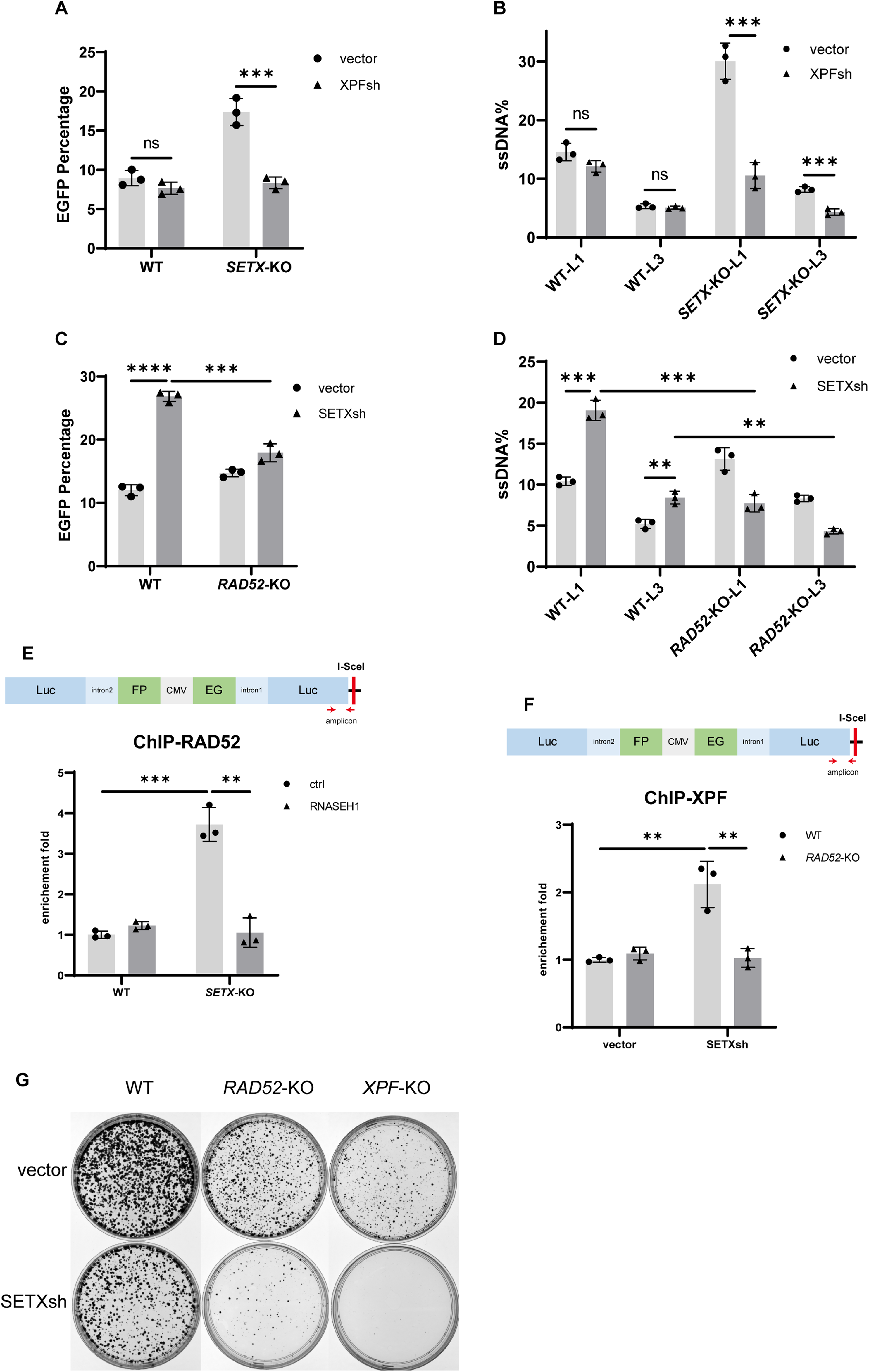
R-loop accumulation activates RAD52-XPF to initiate end resection in SETX-deficient cells. A, B. U2OS WT and *SETX-KO* EGFP-BIR-5085 reporter cells were infected with lentiviruses encoding XPF shRNA or vector. Three days after infection, cells were infected with I-SceI-expressing lentiviruses to induce DSBs. The percentage of EGFP-positive cells was quantified by FACS analysis five days after I-SceI induction (A), and end resection assay was performed two days after I-SceI induction (B). C, D. U2OS WT and *RAD52*-KO EGFP-BIR-5085 reporter cells were infected with lentiviruses encoding SETX shRNA or vector. Three days after infection, cells were infected with I-SceI-expressing lentiviruses to induce DSBs. The percentage of EGFP-positive cells was quantified by FACS analysis five days after I-SceI induction (C), and end resection assay was performed two days after I-SceI induction (D). E. U2OS WT and *SETX-KO* EGFP-BIR-5085 reporter cells, with or without RNASEH1 expression, were infected with I-SceI-expressing lentiviruses to induce DSBs. Two days after infection, cells were harvested for ChIP analysis of RAD52 at the indicated locus on the EGFP-BIR-5085 reporter. F. U2OS WT and *RAD52*-KO EGFP-BIR-5085 reporter cells were infected with lentiviruses encoding SETX shRNA or vector. Three days later, cells were infected with I-SceI-expressing lentiviruses to induce DSBs. Two days after DSB induction, cells were harvested for ChIP analysis of XPF at the indicated locus on the EGFP-BIR-5085 reporter. G. WT, *RAD52*-KO, and *XPF*-KO U2OS cells were infected with lentiviruses encoding SETX shRNA or vector. Three days after infection, cells were resuspended, counted, and 5000 cells were seeded on plates. After two weeks of culture, cells were stained with crystal violet.

Indeed, hyper-BIR and hyper-resection induced by SETX depletion is diminished in *RAD52*-KO cells compared to WT cells (Figure 4C, 4D and S4B), suggesting a role of RAD52 in promoting hyper-resection and hyper-BIR specifically when SETX is deficient. In addition, by ChIP analysis, we observed that RAD52 accumulation at DSBs is significantly increased in the BIR reporter after I-SceI cleavage in *SETX*-KO cells compared to WT cells, which can be suppressed by RNASEH1 overexpression (Figure 4E). We also found that SETX depletion causes XPF accumulation at DSBs generated by I-SceI cleavage in WT cells but not in *RAD52*-KO cells (Figure 4F). These data support the model that RAD52 is associated with R-loops at DSBs and subsequently recruits XPF to cleave R-loops-associated DSB ends, initiating a non-canonical end resection pathway cells (Figure 7A). The involvement of XPF and RAD52 in both hyper-recombination and hyper-resection further supports the idea that hyper-resection in SETX-deficient cells likely contributes to the promotion of BIR.

In the canonical MRN-initiated end resection, BLM, DNA2 and EXO1 are required for further long-range end resection. We depleted BLM, DNA2 and EXO1 and found that they are all indispensable for end resection in both WT and *SETX*-KO cells, as revealed by quantitative end resection assay (Figure 3F, S3C, S3D and S3E) and IR-induced RPA2 phosphorylation (Figure 3G). This suggests that BLM, DNA2 and EXO1 are also involved in the RAD52-XPF hyper-resection pathway, which is activated when R-loops are accumulated at DSBs due to SETX deficiency. We anticipate that bidirectional end resection (Cejka and Symington, 2021) with 3’ to 5’ by the exonuclease activity of MRE11 and 5’ to 3’ by BLM/DNA2 and EXO1 are still employed in the RAD52-XPF hyper-resection pathway (Figure 7A).

As RAD52 and XPF are required for hyper-BIR in SETX-deficient cells, we examined whether RAD52 and XPF would exhibit synthetic lethal interactions with SETX. We found that depletion of SETX in *RAD52*- or *XPF*-KO cells significantly reduces cell viability (Figure 4G and S4C), supporting the critical role of RAD52 and XPF in promoting BIR to repair DSBs with accumulated R-loops at the ends.

### SETX deficiency leads to PCNA ubiquitination, PIF1 loading and BIR onset at DSBs

We showed that SETX loss results in a significant increase in PIF1 recruitment to DSBs (Figure 2C), which may underlie the mechanism driving the onset of BIR at deDSBs in SETX-deficient cells. In yeast, Pif1 interacts with PCNA and this interaction is important for promoting BIR DNA synthesis (Buzovetsky et al., 2017). In mammalian cells, we showed that both PCNA and PIF1 are required for BIR (Li et al., 2021a), and PCNA ubiquitination (PCNA-Ub) is a key step to trigger BIR onset (Shah et al., 2024).

By performing an *in situ* proximity ligation assay (PLA), we showed that in *SETX*-KO cells, the recruitment of both PIF1 (Figure 5A and S5A) and PCNA (Figure 5B) to DSBs, marked by γH2AX after IR, is increased, which is suppressed by RNASEH1 overexpression. Using the antibody that specifically recognizes PCNA-Ub at K164 (Thakar et al., 2020), we observed substantial accumulation of PCNA-Ub at DSBs in *SETX*-KO cells after IR, in contrast to minimal PCNA-Ub accumulation at DSBs in WT cells (Figure 5C). PCNA-Ub at DSBs in *SETX*-KO is suppressed by overexpressing RNASEH1 and inhibiting transcription with 5,6-dichloro-1-beta-D-ribofuranosylbenzimidazole (DRB) (Figure 5C and 5D). These data suggest that accumulated R-loops/hybrids are the trigger to induce the accumulation of PCNA and PIF1 as well as PCNA-Ub at DSBs in SETX-deficient cells.

**Figure 5.**
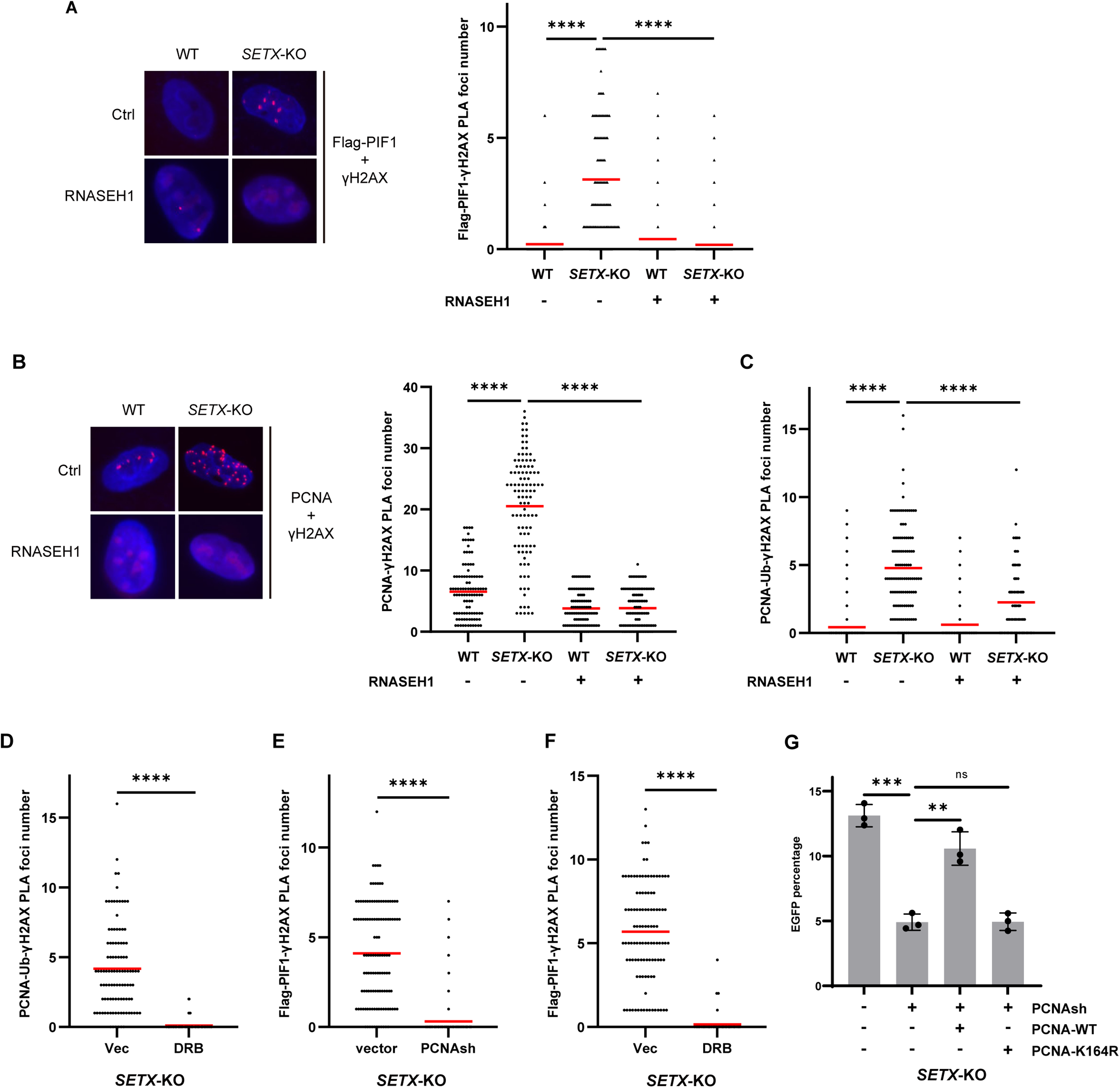
SETX deficiency leads to PCNA ubiquitination, PIF1 loading and BIR onset at DSBs. A. U2OS WT and *SETX*-KO cells expressing Flag-PIF1, with or without RNASEH1 expression, were treated with IR (4 Gy). Two hours later, the co-localization of Flag-PIF1 and γ-H2AX was analyzed by PLA. B, C. U2OS WT and *SETX*-KO cells, with or without RNASEH1 expression, were treated with IR (4 Gy). Two hours later, the co-localization of PCNA (B) or ubiquitinated PCNA (C) with γ-H2AX was analyzed by PLA. D. U2OS *SETX*-KO cells were treated with DMSO or 20 μM DRB for 16 hours prior to IR (4 Gy). Two hours after IR, the co-localization of ubiquitinated PCNA and γ-H2AX was analyzed by PLA. E, F. U2OS *SETX*-KO cells with Flag-PIF1 expression were infected with lentiviruses encoding PCNA shRNA or vector (E), treated with DMSO or 20 μM DRB for 16 hours (F) prior to IR (4 Gy). Two hours after IR, the co-localization of Flag-PIF1 and γ-H2AX was analyzed by PLA. G. U2OS *SETX*-KO EGFP-BIR-5085 reporter cells were infected with lentiviruses to express shRNA-resistant PCNA-WT or PCNA-K164R. Control cells and PCNA-expressing cells were infected with PCNA shRNA or vector. Three days later, all cells were infected with I-SceI-expressing lentiviruses to induce DSBs. The percentage of EGFP-positive cells was quantified by FACS analysis five days after I-SceI induction.

Moreover, PIF1 accumulation at DSBs in *SETX*-KO cells not only can be suppressed by RNASEH1 overexpression (Figure 5A) but also is dependent on PCNA and transcription (Figure 5E, 5F and S5B). These data suggest that PIF1 recruitment to DSBs in SETX-deficient cells is driven by the formation of R-loops/hybrids, with PCNA and PCNA ubiquitination playing a critical role in this recruitment. Additionally, we showed that hyper-BIR in *SETX*-KO cells is diminished after PCNA depletion, which can be recovered by expressing PCNA-WT but not the PCNA ubiquitination mutant PCNA-K164R (Figure 5G and S5C). These findings support the notion that PCNA and PCNA ubiquitination are required for promoting hyper-BIR in SETX-deficient cells, likely through promoting PIF1 recruitment.

### SETX deficiency-induced PCNA ubiquitination at DSBs is dependent on Polα-primase

How PCNA ubiquitination is induced at DSB ends? Typically, PCNA ubiquitination is triggered by replication stalling (Niimi et al., 2008) and is not observed at replication-independent DSBs, such as following IR. We observed PCNA ubiquitination at DSBs in SETX-deficient cells, implying that certain types of replication stalling are induced at DSB ends. Notably, end-fill DNA synthesis has been observed on ssDNA overhangs to antagonize end resection, initiated by Polα-primase that is recruited by shieldin (Mirman et al., 2018; Mirman et al., 2022). Additionally, *de novo* RNA synthesis has also been shown to occur on 3’ ssDNA overhangs (Burger et al., 2019; Gomez-Gonzalez and Aguilera, 2023; Jang et al., 2020; Liu et al., 2021; Michelini et al., 2017; Ohle et al., 2016), and RNA/DNA hybrids accumulate at DSBs in SETX-deficient cells (Cohen et al., 2018). Since PCNA ubiquitination at DSBs in SETX-deficient cells can be suppressed by RNASEH1 overexpression (Figure 5C), we hypothesize that end-fill DNA synthesis would encounter accumulated RNA/DNA hybrids on ssDNA overhangs at DSBs, thereby causing replication stalling to trigger PCNA ubiquitination at DSBs in the absence of SETX (Figure 7B). By performing PLA of PCNA-Ub with γH2AX following IR, we demonstrated that PCNA ubiquitination at DSBs in *SETX*-KO cells is diminished when PRIM1, a subunit of primase, is depleted by shRNA (Figure 6A left and S5D) or when Polα activity is inhibited using the Polα inhibitor CD437 (Han et al., 2016) (Figure 6A right), suggesting that Polα-primase-initiated end-fill DNA synthesis is required for inducing PCNA ubiquitination at DSB ends in SETX-deficient cells, which supports the notion that the conflict of end-fill DNA synthesis with RNA/DNA hybrids on ssDNA overhangs triggers PCNA ubiquitination at DSBs (Figure 7B).

**Figure 6.**
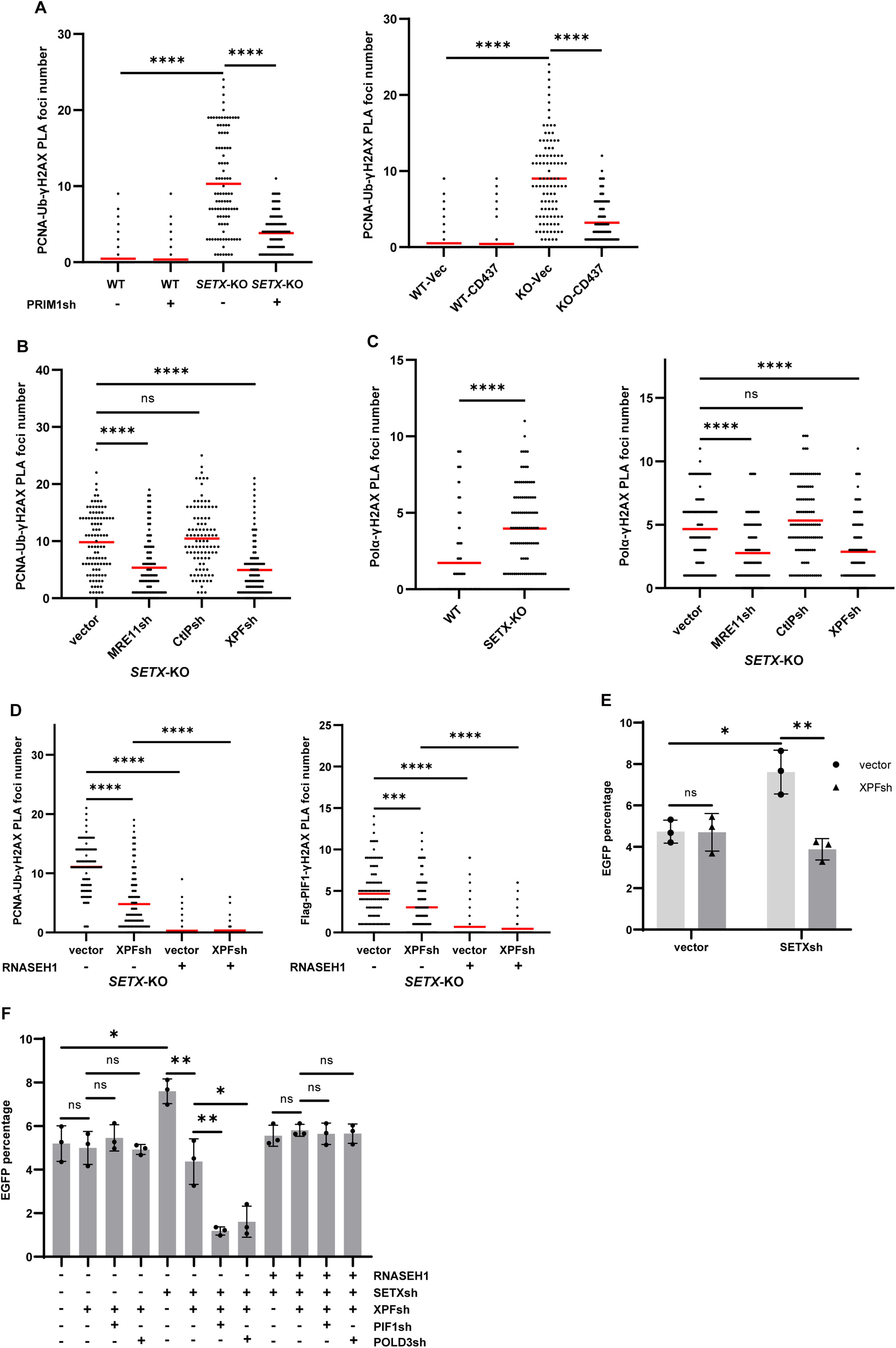
SETX deficiency promotes PCNA ubiquitination through both hyper-resection-dependent and hyper-resection-independent effect. A. U2OS WT and *SETX*-KO cells were infected with PRIM1 shRNA or vector. Three days later, cells were treated with IR (4 Gy) (left). U2OS WT and *SETX*-KO cells were treated with DMSO or CD437 (10 uM) prior to IR (4 Gy) (right). Two hours after IR, the co-localization of ubiquitinated PCNA and γ-H2AX was analyzed by PLA. B. U2OS *SETX*-KO cells were infected with indicated shRNA or vector. Three days later, cells were treated with IR (4 Gy). Two hours after IR, the co-localization of ubiquitinated PCNA and γ-H2AX was analyzed by PLA. C. U2OS WT and *SETX*-KO cells (left), and *SETX*-KO cells infected with indicated shRNAs (right) were treated with IR (4 Gy). Two hours later, the co-localization of Polα and γ-H2AX was analyzed by PLA. D. U2OS *SETX*-KO cells expressing Flag-PIF1, with or without RNASEH1 expression, were infected with XPF shRNA or vector. Three days later, all cells were treated with IR (4 Gy). Two hours after IR, the co-localization of ubiquitinated PCNA and γ-H2AX (left), and the co-localization of Flag-PIF1 and γ-H2AX (right), were analyzed by PLA. E. U2OS EGFP-HR-5633 reporter cells were infected with vector, XPF shRNA, SETX shRNA or both XPF shRNA and SETX shRNA. Three days later, cells were infected with lentiviruses expressing I-SceI to induce DSBs. The percentage of EGFP-positive cells was quantified by FACS analysis five days after DSB induction. F. U2OS EGFP-HR-5633 reporter cells with or without RNASEH1 expression were infected with indicated shRNAs. Three days later, cells were infected with lentiviruses expressing I-SceI to induce DSBs. The percentage of EGFP-positive cells was quantified by FACS analysis five days after DSB induction.

**Figure 7.**
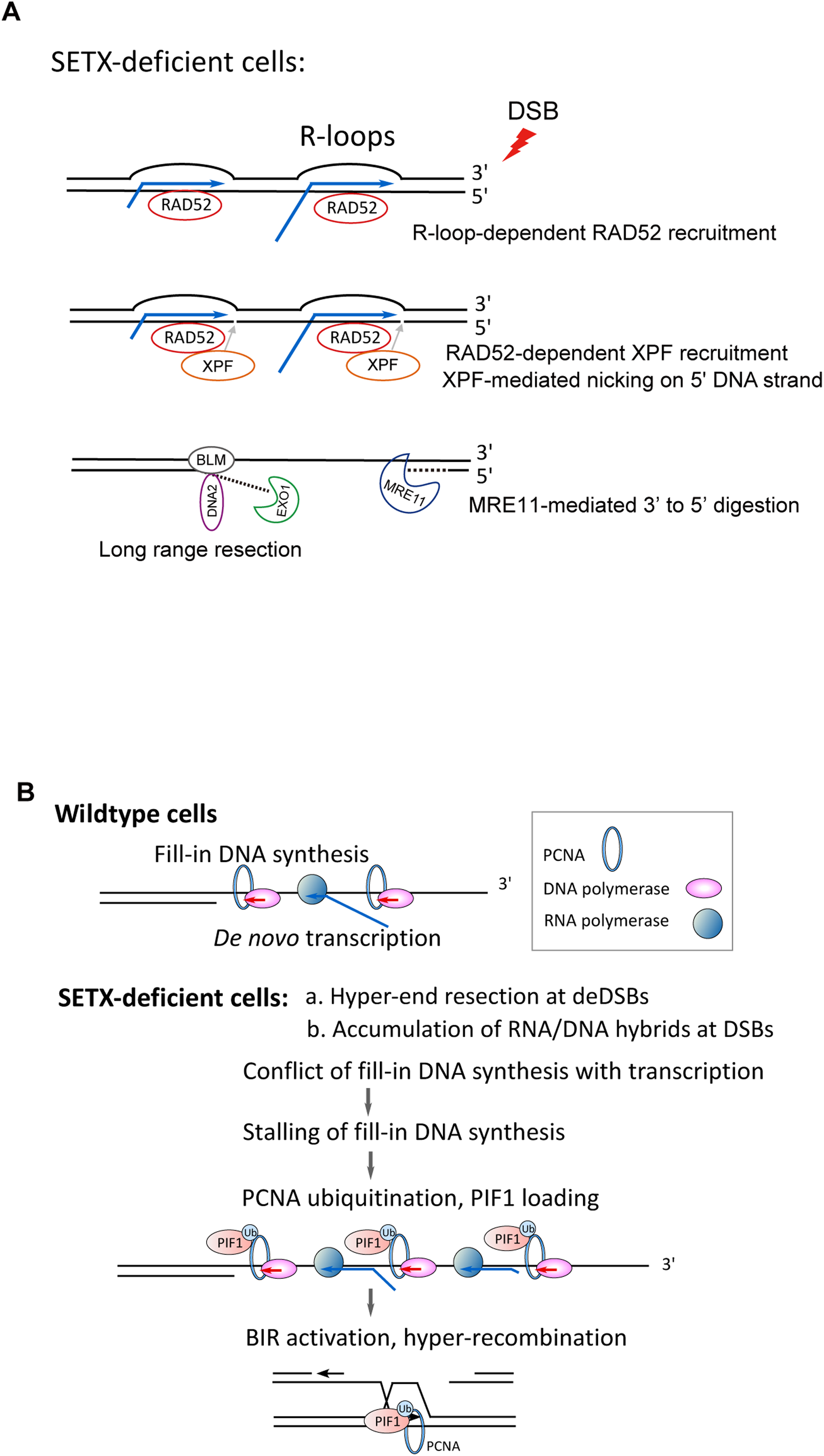
Working Models for the induction of hyper-end resection and BIR onset at deDSBs in SETX-deficient cells. A. Illustration of non-canonical end resection pathway in SETX-deficient cells. SETX deficiency causes more R-loop formation on DSB ends and RAD52 recruitment, which in turn recruits XPF. Instead of MRE11, XPF generates nicks on the 5’ strand of DSBs with accumulated R-loops, followed by removal of the remaining 5’ strand towards the DSB ends by MRE11 through its 3’ to 5’ exonuclease activity, and by long-range resection of 5’ strand away from the DSB ends by EXO1 and BLM/DNA2. B. Working model of BIR activation at deDSB ends. After end resection at DSBs, while Polα-primase-initiated end-fill occurs at ssDNA overhangs, *de novo* RNA synthesis also takes place. In WT cells, with the action of SETX and possibly other R-loop/hybrid resolvases, *de novo* RNA synthesis only generates very limited RNA/DNA hybrids on the ssDNA overhangs, which do not interfere with end-fill DNA synthesis to induce PCNA ubiquitination (top). However, in SETX-deficient cells, RNA/DNA hybrids on ssDNA overhangs are accumulated, which stall end-fill DNA synthesis, thereby promoting PCNA ubiquitination, PIF1 recruitment, and subsequent BIR activation (bottom). Additionally, hyper-end resection due to SETX deficiency allows more loading of end-fill DNA synthesis replicons on resected DSB ends, leading to stronger PCNA ubiquitination and PIF1 loading, which contributes to the hyper-recombination observed in SETX-deficient cells.

### Hyper-end resection induced by SETX deficiency promotes but is not required for PCNA ubiquitination and BIR onset at DSBs

We observed that hyper-BIR is correlated with hyper-resection when SETX is deficient. We also found that in SETX-deficient cells, XPF depletion suppresses both hyper-resection and hyper-BIR, whereas CtIP inactivation is dispensable for both processes. This raises the question of whether hyper-resection observed in SETX-deficient cells is critical for promoting BIR by driving PCNA ubiquitination and PIF1 recruitment at DSBs. To test this, we first determined whether hyper-resection is required for PCNA ubiquitination observed in the absence SETX. We depleted XPF, MRE11 or CtIP in SETX-deficient cells and found that depleting XPF or MRE11 reduces PCNA ubiquitination; however, a substantial level of PCNA ubiquitination persists, whereas CtIP depletion has no effect (Figure 6B and S5E). Since SETX deficiency-induced hyper-resection depends on XPF and MRE11 but not CtIP, these findings suggest that hyper-resection facilitates PCNA ubiquitination at DSBs. However, the presence of substantial PCNA ubiquitination even in the absence of hyper-resection suggests that while hyper-resection enhances PCNA ubiquitination, it is not essential for triggering this process.

It is possible that hyper-resection in SETX-deficient cells generates longer ssDNA overhangs, allowing the assembly of more end-fill DNA synthesis replicons, which in turn produces stronger replication stalling signals to induce PCNA ubiquitination compared to WT cells (Figure 7B). In support of this, we showed that Polα recruitment to DSBs is increased in *SETX*-KO cells compared to WT cells, which can be suppressed by depleting MRE11 or XPF but not CtIP (Figure 6C), suggesting that hyper-resection indeed triggers increased loading of Polα-primase to establish additional fill-in synthesis replicons on ssDNA overhangs at DSBs when SETX is deficient.

Notably, depleting XPF in *SETX*-KO cells reduces end resection to the level comparable to WT cells (Figure 5B) and concurrently abolishes hyper-BIR in *SETX*-KO cells (Figure 5A), but substantial PCNA ubiquitination (Figure 6B and 6D left) and PIF1 loading to DSBs (Figure 6D right) persist although reduced. We overexpressed RNASEH1 to reduce R-loops/hybrids accumulation and found that RNASEH1 overexpression reduces PCNA ubiquitination and PIF1 loading to DSBs in *SETX*-KO cells expressing XPF shRNA (Figure 6D). This suggests that even in the absence of hyper-resection, RNA/DNA hybrids accumulated at DSBs are sufficient to induce PCNA ubiquitination and PIF1 loading to DSBs.

To test whether PCNA ubiquitination remaining in *SETX*-KO cells with XPF shRNA expression would still induce the onset of BIR mechanism at DSBs, we analyzed whether HR/STGC assayed by U2OS (EGFP-HR/STGC-5633) reporter cell line is dependent on PIF1 and POLD3. We showed that hyper-recombination (HR) is reduced in SETX-depleted cells after depleting XPF, to the level comparable to that in WT (Figure 6E and Figure 6F, compare lane 5 and 6), consistent with the notion that hyper-recombination is dependent on XPF-driven hyper-resection. However, in contrast to cells expressing XPF shRNA alone, HR/STGC in cells co-expressing shRNAs for both SETX and XPF shows dependence on PIF1 and POLD3 (Figure 6F, compare lane 3 and 4 with lane 7 and 8, Figure S5F). Furthermore, RNASEH1 overexpression abolishes the dependence of HR/STGC on PIF1 and POLD3 in cells co-depleted for SETX and XPF (Figure 6F), suggesting that the BIR mode, which requires PIF1 and POLD3, is still established at DSBs in SETX- and XPF-deficient cells. Collectively, these data suggest that accumulated RNA/DNA hybrids at DSBs in SETX-deficient cells induce PCNA ubiquitination and PIF1 loading even in the absence of XPF-driven hyper-resection, which triggers the establishment of using the BIR mode at DSBs. Hyper-resection promotes hyperrecombination using the BIR mode which has already been established at deDSBs in SETX-deficient cells.

## Discussion

In this report, we showed that SETX deficiency leads to hyper-recombination, which is promoted by the activation of the BIR mechanism. Our working model is that in the absence of SETX, accumulated RNA/DNA hybrids on the ssDNA overhangs at DSBs conflict with end-fill DNA synthesis initiated by Polα-primase, which induces PCNA ubiquitination and PIF1 loading at DSBs, leading to the onset of BIR (Figure 7B). Additionally, accumulation of R-loops at DSBs prior to end resection in SETX-deficient cells triggers XPF-dependent non-canonical hyper-resection (Figure 7A), leading to the assembly of more end-fill DNA synthesis replicons on the elongated ssDNA overhangs, which in turn induces BIR-mediated hyper-recombination. These studies provide new insights into BIR activation and highlight the importance of R-loop homeostasis at DSBs in the HR and BIR repair pathway selection.

### PCNA ubiquitination and PIF1 recruitment for BIR activation at DSBs

It is well-accepted that BIR is primarily used to repair seDSBs, such as at broken forks and eroding telomeres (Kramara et al., 2018; Wu and Malkova, 2021b). Our previous study showed that PCNA is accumulated and ubiquitinated at seDSBs on broken forks, recruiting PIF1 to activate the BIR mechanism for mediating both BIR/LTGC and HR/STGC (Li et al., 2021a; Shah et al., 2024) (Figure S1B left). At deDSBs, PCNA ubiquitination and PIF1 loading are minimal, in sharp contrast to high levels of PCNA ubiquitination and PIF1 loading at seDSBs upon fork breakage (Figure S1B right). We showed that 53BP1 deficiency induces BIR onset at DSBs by triggering increased PCNA loading and ubiquitination (Shah et al., 2024). Therefore, PCNA accumulation and ubiquitination are the key events for establishing the BIR mode.

The major difference in HR/STGC and BIR is the DNA synthesis length during D-loop migration. HR replisomes and BIR replisomes share common components such as Polδ but are different in at least some components. Incorporating PIF1 and other unknown factors likely enhances the potency and processivity of BIR replisomes for DNA replication. We propose that the assembly of BIR replisomes at DSBs triggers a switch from using the HR mechanism to the BIR mechanism (Li et al., 2021a). In contrast to HR replisomes, which are exclusively used for HR/STGC, BIR replisomes can be used for both HR/STGC and BIR/LTGC. Thus, HR/STGC would still be detected in the EGFP-HR/STGC reporter assay, even if the repair mechanism at DSBs has switched to using the BIR pathway, showing the dependence on PIF1 and POLD3.

### Establishing the BIR mechanism at deDSBs accumulated with RNA/DNA hybrids in SETX-deficient cells

Importantly, the hyper-recombination that we observed in SETX-deficient cells is not merely an increase of recombination activity but involves a shift of pathway usage; upon loss of SETX, elevated HR/STGC requires PIF1, showing BIR characteristics (Figure 2B). We observed that increased PCNA ubiquitination at DSBs in SETX-deficient cells is suppressed by RNASEH1 overexpression and transcription inhibition, suggesting that transcription and possibly accumulated RNA/DNA hybrids trigger PCNA ubiquitination at DSBs. *De novo* RNA synthesis has been shown to occur on the 3’ ssDNA overhangs at DSBs (Burger et al., 2019; Gomez-Gonzalez and Aguilera, 2023; Jang et al., 2020; Liu et al., 2021; Michelini et al., 2017; Ohle et al., 2016), and we speculate that accumulated RNA/DNA hybrids cause conflict with end-fill DNA synthesis on ssDNA overhangs in SETX-deficient cells. This is supported by the observation that inhibition of Polα-primase-dependent end-fill DNA synthesis suppresses PCNA ubiquitination and PIF1 recruitment to activate BIR. This mechanism mirrors the induction of PCNA ubiquitination and BIR activation at DSBs in 53BP1-deficient cells, where end-fill DNA synthesis, initiated by shieldin-independent loading of Polα-primase, is stalled on ssDNA overhangs (Shah et al., 2024). We propose that while various causes, such as replication fork collapse, accumulation of RNA/DNA hybrids at deDSBs or 53BP1 deficiency, interfere with DNA replication or synthesis, the resulting PCNA ubiquitination—whether at broken replication forks or at DSB ends—would serve as a common signal for PIF1 recruitment and the activation of the BIR mechanism.

We further propose that abnormally accumulated RNA/DNA hybrids at deDSBs are one of the determinants for the repair pathway selection favoring BIR/LTGC. In WT cells, under normal situations, R-loops/hybrids are present at deDSBs often in the transcriptionally active regions but with an appropriate level, and transcription-associated homologous recombination repair (TA-HRR) can be engaged to promote HR/STGC (Yasuhara et al., 2018). However, uncontrolled accumulation of R-loops/hybrids at deDSBs could be one of the signals to induce BIR onset at deDSBs. We showed that in SETX-deficient cells, R-loops/hybrids are abnormally accumulated, which induces PCNA ubiquitination and triggers the onset of BIR at deDSBs. Therefore, in addition to repairing seDSBs at collapsed forks and broken telomeres, BIR is also involved at deDSBs in the transcriptionally active regions when R-loops/hybrids are abnormally accumulated due to disruptions in R-loop/hybrid homeostasis at DSB ends. Notably, it has been shown that telomeric R-loop formation caused by non-coding RNA TERRA promotes telomere maintenance using the BIR mechanism (In et al., 2025; Yadav et al., 2022). Furthermore, MiDAS, using BIR mechanism, often occurs at common fragile sites (CFSs) (Bhowmick et al., 2023), where R-loops tend to accumulate due to TRC at large genes (Helmrich et al., 2011). Accumulated R-loops at CFSs may signal the onset of BIR to reduce the use of end joining, which would generate deletions, thereby safeguarding genome integrity by using BIR to complete replication of the unreplicated DNA regions (Figure S6). In this context, it has been shown that loss of SETX leads to increased MiDAS (Said et al., 2022).

### The mechanism underlying hyper-resection in SETX-deficient cells

We observed that in SETX-deficient cells, end resection is hyperactive, mediated by a non-canonical end resection pathway that is independent of the endonuclease activity of MRE11 and CtIP, but requires RAD52 and XPF. In one scenario, persistence of R-loops at DSBs may inhibit the MRE11 nuclease activity, preventing the endonucleolytic cleavage to initiate end resection, consistent with the *in vitro* assay that RNA-DNA hybrids inhibit the MRE11 nuclease activity (Chang et al., 2019). Alternatively, XPF, recruited by RAD52 to R-loops at DSBs, may override the function of MRE11 and CtIP in endonucleolytic cleavage by creating nicks more internally on the 5’ strands (Figure 7A). XPF has been implicated in processing R-loops to generate DSBs (Makharashvili et al., 2018; Sollier et al., 2014; Stork et al., 2016).

Consistent with our findings, hyper-resection was also observed in yeast Sen1-deficient cells, (Rawal et al., 2020). However, in a previous study using SETX depletion in mammalian cells, hyper-resection was not detected despite the accumulation of R-loops/hybrids at DSB ends (Cohen et al., 2018), and this discrepancy could be related to the level of SETX depletion. In the yeast Sen1-deficient model, Mre11 endonuclease is proposed to create a nick to initiate end resection, followed by Dna2 cleaving long 5’ flaps of ssDNA in R-loops to promote extensive end resection, while Sgs1 (BLM homologue) and Exo1 are dispensable (Rawal et al., 2020). We found that in SETX-deficient cells, hyper-resection occurs independently of the endonuclease activity of MRE11 and CtIP, suggesting differences in the underlying mechanism. We propose that when SETX is deficient, RAD52 recognizes accumulated R-loops at DSBs, and recruits XPF to cleave the 5’ side of R-loops on the 5’ strands of DSBs to initiate end resection (Figure 7A). With anticipated multiple transcription forks at transcriptional active DSBs, XPF could make a series of cleavages on the 5’ strand of a DSB, thereby robustly accelerating end resection and resulting in hyperactive end resection phenotype. The exonuclease activity of MRE11, as well as BLM/DNA2 and EXO1, are still required for bidirectional processing to complete end resection.

Studies on the roles of R-loops at deDSBs in end resection by disrupting the functions of different R-loop resolvases yield quite diverse results, including impaired end resection (Alfano et al., 2019; Matsui et al., 2020; Ohle et al., 2016; Sessa et al., 2021; Yu et al., 2020), no discernible influence on end resection (Li et al., 2016), or accelerated end resection [(Domingo-Prim et al., 2019; Rawal et al., 2020), this study]. This is likely attributed to the distinct functions of different R-loop resolvases, with their impaired activities causing variations in the structure, extent, and level of R-loops/hybrids formed at deDSBs. We showed that R-loops accumulated at DSBs in SETX-deficient cells can be recognized by RAD52-XPF, triggering a non-canonical hyper-resection mechanism which overrides the requirement of MRE11 endonuclease activity and CtIP. However, accumulated R-loops or hybrids due to defects in other hybrid resolvases may obstruct the enzymatic activities of MRE11, BLM, or EXO1, as suggested by *in vitro* experiments (Chang et al., 2019; Daley et al., 2020), hindering end resection.

### SETX deficiency results in hyper-recombination

SETX deficiency does not only promote the onset of BIR but also induces hyper-recombination using the BIR mode for both HR/STGC and BIR/LTGC. Inactivation of XPF and MRE11 prevents both hyper-resection and hyper-recombination, suggesting that hyper-resection facilitates hyper-recombination. We anticipate that while the conflict between accumulated RNA/DNA hybrids and end-fill DNA synthesis at ssDNA overhangs serves as the essential signal to trigger PCNA ubiquitination, PIF1 loading, and BIR onset at deDSBs in SETX-deficient cells, over-resected ends further induce hyper-recombination activity via the BIR mechanism. Elongated ssDNA overhangs could harbor more stalled end-fill replicons, potentially triggering stronger PCNA ubiquitination and BIR activation (Figure 7B), contributing to hyper-recombination. Additionally, it remains possible that these longer ssDNA overhangs may provide extended homology to stabilize D-loops, thereby promoting recombination.

While hyper-recombination is induced in SETX-deficient cells, HR defects have also been reported when other RNA resolvases are defective (Alfano et al., 2019; D’Alessandro et al., 2018; Matsui et al., 2020; Ohle et al., 2016; Sessa et al., 2021; Yu et al., 2020). In yeast, both depletion of Rnh1 and Rnh201 and overexpression of Rnh1 result in HR defects (Ohle et al., 2016), suggesting a delicate balance in R-loops/hybrid formation for regulating HR. Since both formation and resolution of R-loops/hybrids at DSBs are crucial for HR, different hybrid resolvases may function at distinct stages and could positively or negatively influence HR. We speculate that hybrids accumulated on the 3’ ssDNA overhangs at deDSBs in SETX-deficient cells are removed subsequently by other resolvases, allowing hyper-HR and hyper-BIR in the absence of SETX.

### Connection to cancer treatment

SETX gene is frequently mutated in cancers, and its expression is often downregulated in a variety of cancers, akin to other tumor suppressor genes (Rhodes et al., 2007). We uncovered a DSB pathway shift accompanied with hyper-recombination at deDSBs from using the HR mechanism to the BIR mechanism in SETX-deficient cells and demonstrated synthetic lethal interactions of SETX with PIF1, RAD52 and XPF. The pronounced reliance of SETX-deficient cells on PIF1, RAD52 and XPF for cell viability lays groundwork for establishing new treatment strategies through inhibiting the hyperactive BIR pathway to selectively eradicate cancers with compromised SETX function, while preserving viability of normal cells.

## Resource availability

### Lead contact

Further information and requests for resources and reagents should be directed to and will be fulfilled by the lead contact, Xiaohua Wu.

### Materials availability

This study did not generate new unique reagents.

### Data and code availability

The data reported in the paper will be shared upon request to the lead contact.

This study did not generate original code.

Any additional information required to reanalyze the data reported in this paper is available from the lead contact upon request.

## Materials and Methods

### Cell cultures

U2OS (ATCC), 293T (ATCC) and U2OS cells with *RAD52*-KO (Liu et al., 2023) cells were cultured in DMEM media supplemented with 10 % fetal bovine serum (FBS) and 1 % penicillin/streptomycin at 37°C, 5% CO2.

### Plasmid construction

The N-terminal Flag-tagged WT and helicase-dead (P-loopΔ) SETX-expressing plasmids are gifts from Dr. Craig Bennett (Bennett et al., 2020). Their coding sequences were cloned into pCDH-CMV-MCS-EF1-puro lentiviral vector (System Biosciences). The C-terminal V5-tagged WT RNASEH1 plasmids are gifts from Dr. Xiangdong Fu (Chen et al., 2019). Their coding sequences were cloned into pCDH-CMV-MCS-EF1-puro lentiviral vector (System Biosciences).

### Generation of knock out (KO) cell line by CRISPR

Two sgRNAs sequences targeting coding sequence of specific genes were cloned into sgRNA/Cas9-mCherry vector derived from pSpCas9(BB)-2A-Puro (PX459) V2.0 vector (Addgene #62988). The two sgRNA plasmids were transfected together into target cell lines using standard PEI protocol. Two days later, transfected cells were sorted for mCherry-positive events and plated for single clones.

These single clones were screened with PCR or Western blotting.

sgRNA sequences for *SETX*-KO:

aggctaaatccgttgatcac and TGTTGAAGCACTTTGTCGGA

sgRNA sequences for *XPF*-KO:

GGGCTAGTAGTGTGCGCCCG and GCTGGTGCTCAACACGCAGC

### shRNA interference

Oligoes containing shRNA target sequences were cloned into pLKO.1-blast lentiviral vector (Addgene #26655). shRNA sequences used in this study are listed below.

SETXsh CGGCAGAAGGATTGTGTTATT

MRE11sh GATGAGAACTCTTGGTTTATT

CtIPsh GAGCAGACCTTTCTCAGTATA

BLMsh GAGCACATCTGTAAATTAATT

DNA2sh AGTTTGTGATGGGCAATATTT

EXO1sh ACTCGGATCTCCTAGCTTTTG

XPFsh AAGACGAGCTCACGAGTATTC

PIF1sh GAAGACAGGTGCTCCGGAAGC

PCNAsh GTGGAGAACTTGGAAATGGAA

PRIM1sh AGCATCGTCTCTGGGTATATT

Knockdown efficiency of individual shRNA was confirmed by Western blotting of whole-cell lysate or RT-qPCR with primers listed below.

EXO1_quanti_F TCGGATCTCCTAGCTTTTGGCTG

EXO1_quanti_R AGCTGTCTGCACATTCCTAGCC

DNA2_quanti_F GATTTCTGGCACCAGCATAGCC

DNA2_quanti_R ACACCTCATGGAGAACCGTACC

CtIP_quanti_F TGGCAGACAGTTTCTCCCAAGC

CtIP_quanti_R GGCTCCACAAACGCTTTCTGCT

PIF1_quanti_F CATCCACAAGAGCCAAGGCAT

PIF1_quanti_R GGTGGCATAGAAGTGCAGCA

POLD3_quanti_F TTCTGTCAAGAGCTCAAGTGG

POLD3_quanti_R GTGCAGGATTCACTCTCGTAG

BLM_quanti_F GGTGATAAGACTGACTCAGAAGC

BLM_quanti_R AACGTGCCAAGAGCTTCCTCTC

GAPDH_quanti_F GTCTCCTCTGACTTCAACAGCG

GAPDH_quanti_R ACCACCCTGTTGCTGTAGCCAA

PCNA_quanti_F CCTGCTGGGATATTAGCTCCA

PCNA_quanti_R CAGCGGTAGGTGTCGAAGC

XPF_quanti_F CGCAGAGCTAACCTTTGTTCGG

XPF_quanti_R TCCGCAAAGCAGTGAGATAGCG

MRE11_quanti_F CAGCAACCAACAAAGGAAGAGGC

MRE11_quanti_R GAGTTCCTGCTACGGGTAGAAG

### Colony formation assay

Colony formation assay was carried out to examine the potential synthetic lethality effect between SETX and other tested genes. Cells were infected with lentivirus to knock down indicated genes. Three days after infection, cells were resuspended and counted. For each condition, 5000 cells were plated on 10cm plate. Two weeks later, plates with growing colonies were stained with 0.5 % crystal violet in 20 % methanol.

### Western blotting

Cells cultured in 6-well plate were lysed in RIPA buffer (50 mM Tris–HCl, pH 8.0, 150 mM NaCl, 5 mM EDTA, 1 % NP-40, 0.5 % sodium deoxycholate, and 0.1 % SDS). Fresh lysates were supplemented with eqaul amount of 2X loading buffer (100 mM Tris–Cl, pH 6.8, 4 % SDS, 0.2 % bromophenol blue, 20 % glycerol, 200 mM DTT) and then boiled at 95°C for 5 min. Next, sampels were separated on 6–15 % SDS–PAGE and proteins were transferred to nitrocellulose membrane (1620115, Bio-Rad) using the Trans-Blot Turbo Transfer System from Bio-Rad. Antibodies used were followed: SETX (NB100-57542, Novus Biologicals), KU70 (sc-17789, Santa Cruz Biotechnology), FLAG (F1804, Sigma-Aldrich), V5 (R96025, Invitrogen), XPF (sc-136153, Santa Cruz Biotechnology), RPA32 (Ser4/Ser8) (A300-245A, Bethyl Laboratories), γH2AX (2577, Cell Signaling Technology), PCNA (sc-56, Santa Cruz), GAPDH (AC002, ABclonal), RAD52 (A5186, ABclonal), PRIM1 (10773-1-AP, Proteintech), HA.11 (MMS-101P, COVANCE).

### Immunostaining

Cells were seeded on coverslips in 6-well plate and cultured overnight. The next day, cells were first pre-extracted with 0.5 % Trion X-100 in cold PBS for 5mins, then fixed with 2 % formaldehyde for 10 mins at room temperature. Fixed cells were blocked with 5 % milk in PBS for 30 mins and then incubated overnight with indicated primary antibodies diluted in PBS at 4°C. The following day, cells were washed with PBS for three times and further incubated with secondary antibody and DAPI. Finally, slides were sealed with mounting media (P36930, Invitrogen) to prevent quenching. Images were taken on Nikon ECLIPSE Ni-L microscope. S9.6 antibody (ENH001, Kerafast) was used.

### In situ proximity ligation assay (PLA)

Cells were seeded on coverslips in 6-well plate and cultured overnight. The next day, cells were treated with 4 Gy IR. Two hours later, cells were fixed and blocked the same way as immunostaining and then incubated with primary antibody overnight at 4°C. The following day, PLA was performed using the Duolink PLA kit (DUO92001 and DUO92005, Sigma-Aldrich) following the manufacturer’s protocol. Finally, slides were sealed with mounting media (P36930, Invitrogen) to prevent quenching. Images were taken on Nikon ECLIPSE Ni-L microscope. Primary antibodies used were as followed: FLAG (F1804, Sigma-Aldrich), PCNA (SC-56, Santa Cruz), γH2AX (07164, Upstate) and PCNA-Ub (13439, Cell Signaling Technology), DNA polymerase α (sc-137021, Santa Cruz).

### Chromatin immunoprecipitation (ChIP)

Cells growing on 10 cm plates were fixed with 1 % formaldehyde for 10 min at room temperature and glycine was added to the final concentration of 0.125 M. Cells were scraped from the plates and lysed with 0.5 % NP-40 buffer (10 mM Tris-HCl, 10 mM NaCl, 0.5 % NP-40, freshly added Roche cOmplete protease inhibitor cocktail) for 15mins at 4°C. Whole lysates were sonicated with Bioruptor for 15 cycles (30s on/30s off) and then centrifuged to remove debris. Supernatent was collected and incubated with Protein G dynabeads which have been pre-loaded with indicated antibody, for 4 hours at 4°C. After IP, beads were washed three times with 0.5 % NP-40 buffer and once with TE buffer. Finally, beads was incubated in elution buffer (10 mM Tris-HCl, 1.0 % SDS, 1.0 mM EDTA) overnight at 65°C to reverse the crosslinking. The next day, IP products and input were subjected to DNA extraction using phenol-chloroform. Recovered DNA was checked by qPCR to calculate the relative enrichement of target protein on specific locus. Primers used for ChIP were listed below.

At I-SceI-induced DSB end:

ChIP_L1F GAATCCATCTTGCTCCAACACCCC

ChIP_L1R CACAAACACAACTCCTCCGCGC

Antibodies used were as followed: S9.6 (ENH001, Kerafast), RAD52 (A5186, ABclonal), XPF (sc-136153, Santa Cruz Biotechnology).

### End resection quantification assay

The end resection assay in reporter cells was adopted from publications of Dr. Tanya Paull’s lab (Zhou et al., 2014) and modified according to our BIR reporter sequence. The BIR reporter cells were infected with lentivirus expressing I-SceI to induce DSBs. Two days after infection, cells were harvested for genomic DNA extraction using Puregene Kits from QIAGEN. 1 ug genomic DNA was digested separately with NEB restriction enzyme BsiHKAI (left DSB end), ApoI (right DSB end) or SacII (mock) overnight. Restriction enzyme digested genomic DNA was used as template for qPCR with indicated primer listed below.

L1:

End-Resection_L1F GCTCCAACACCCCAACATCTTCGAC

End-Resection_L1R CGGTACTTCGTCCACAAACACAACTCC

L3:

End-Resection_L3F GGTGAACTTCAAGATCCGCCACAAC

End-Resection_L3R CAGCTCGTCCATGCCGAGAG

R1:

End-Resection_R1F gaactagtggatcCCCTACCGTTCG

End-Resection_R1R ggtggatgtggaatgtgtgcgagg

R3:

End-Resection_R3F CGAGATCAGCAGCCTCTGTTCCACA

End-Resection_R3R CTGTCACCAATCCTGTCCCTAGTGCA

### Statistical analysis

Student’s t-test was run in two-tailed non-paired mode to examine the significance of differences between samples. In all experiments, error bars show standard deviation (SD) of at least three independent experiments. The P value is labeled as * P < 0.05, ** P < 0.01, *** P < 0.001, **** P < 0.0001, and ns (not significant) P > 0.05.

## Acknowledgement

We would like thank Dr. Craig Bennett (University of Lincoln, UK) for providing the cDNA of WT and helicase-dead (P-loopΔ) mutant SETX, Dr. Xiangdong Fu (UCSD) for providing the WT RNASEH1 constructs. Plasmids pLKO.1-blast vector (26655) and pSpCas9(BB)-2A-Puro (PX459) V2.0 (62988) are from Addgene. This study is funded by NIH grants CA244912, GM141868 and CA187052 to X.W.

## Author contributions

T.W. designed and performed experiments and analyzed the data; Y.L. and L.S. conducted some experiments. X.W. is responsible for the overall project’s planning and experimental design. T.W. and X.W. wrote the manuscript.

---

**Fig S1 related to Fig 1.**
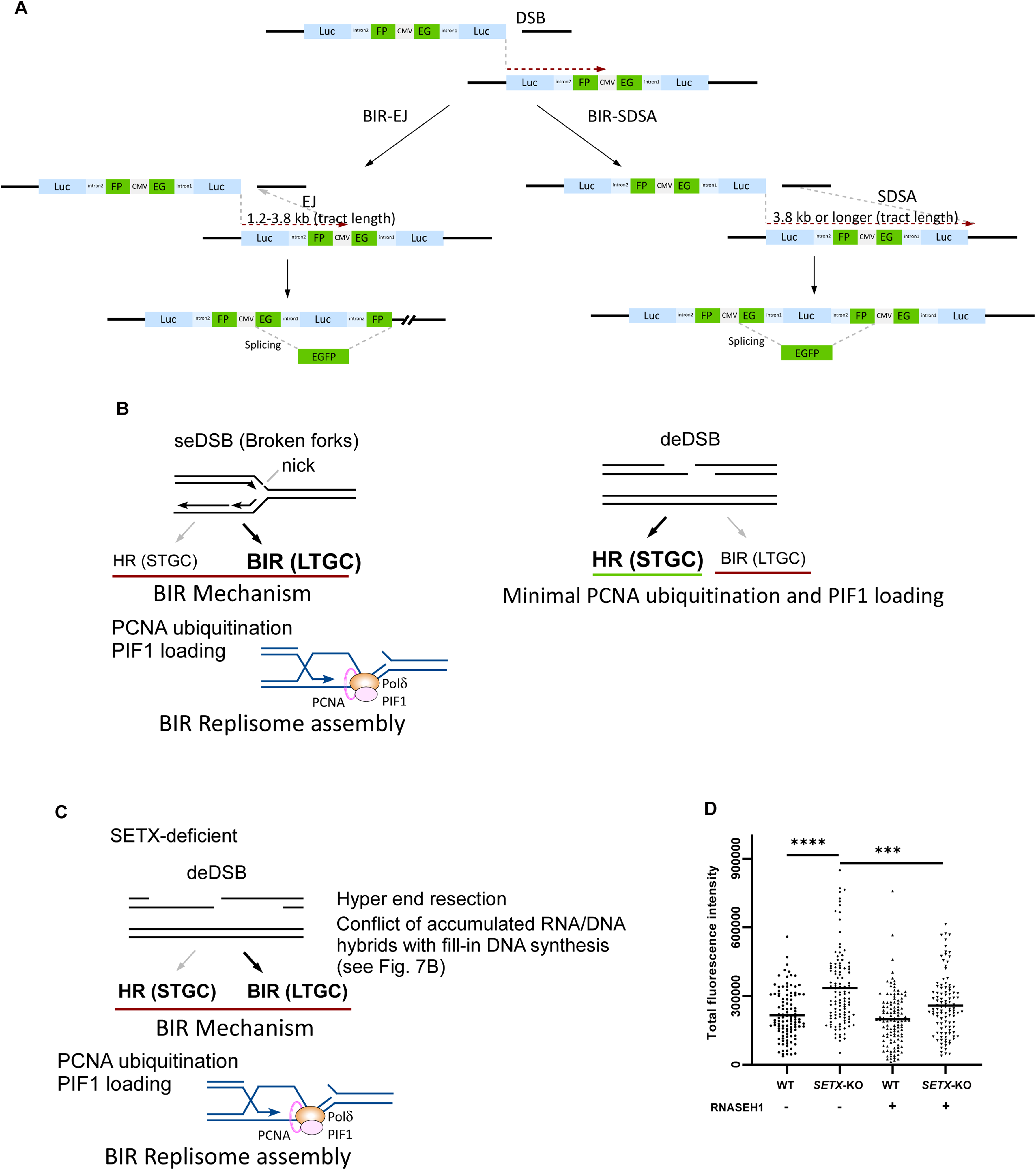
DSB repair pathway choice. A. Schematic drawing of repair products (BIR-EJ or BIR-SDSA) in EGFP-BIR-5085 reporter. B. BIR activation at seDSB on broken forks (left) and at deDSB (right). C. BIR activation at deDSB in SETX-deficient cells. D. U2OS WT and SETX-KO EGFP-BIR-5085 reporter cells were infected with lentiviruses expressing RNASEH1. Control cells and RNASEH1-expressing cells were fixed and subjected to immunostaining with S9.6 antibody to quantify R-loop levels.

**Fig S2 related to Fig 2.**
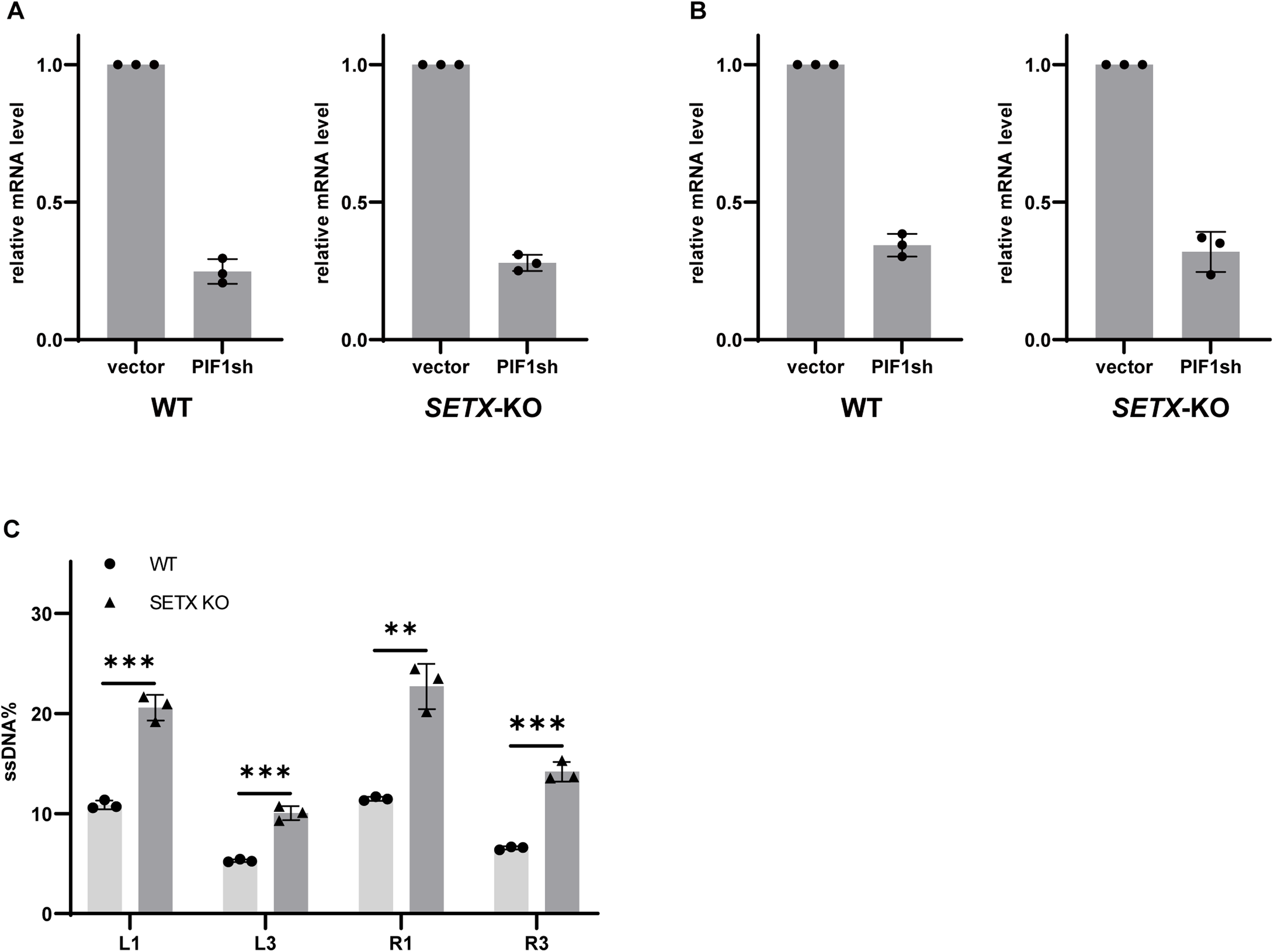
Determining the dependence on essential BIR genes after SETX inactivation and determining the effect of R-loop presence on end resection assay.

**Fig S3 related to Fig 3.**
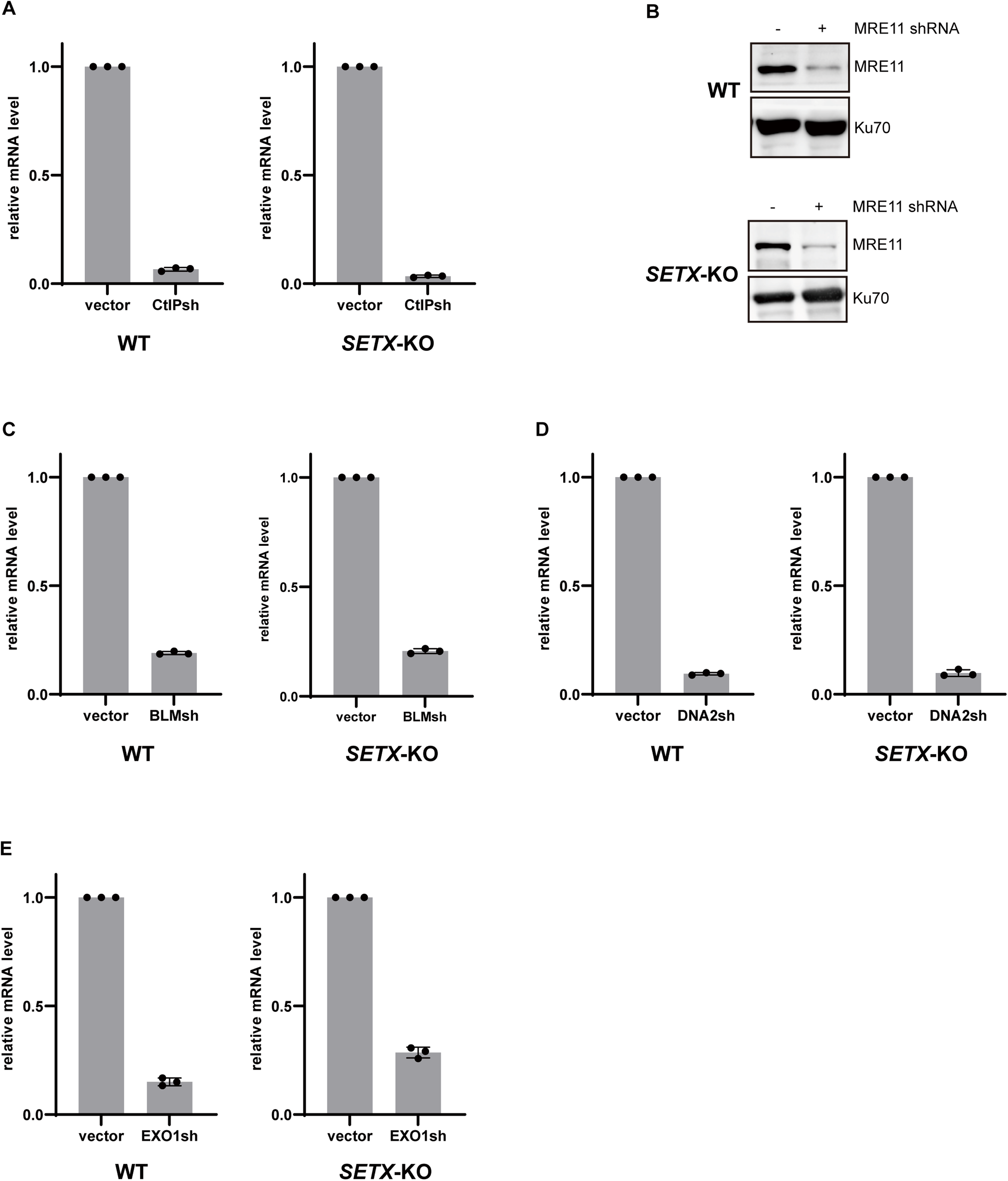
Determining the dependence on canonical end resection pathway in SETX-deficient cells. A. qPCR results showing the efficiency of CtIP shRNA in WT and *SETX*-KO U2OS EGFP-BIR-5085 reporter cells. B. Western blotting showing the efficiency of MRE11 shRNA in WT and *SETX*-KO U2OS EGFP-BIR-5085 reporter cells. C,D,E.qPCR results showing the efficiency of BLM shRNA, DNA2 shRNA, and EXO1 shRNA in WT and *SETX*-KO U2OS EGFP-BIR-5085 reporter cells.

**Fig S4 related to Fig 4.**
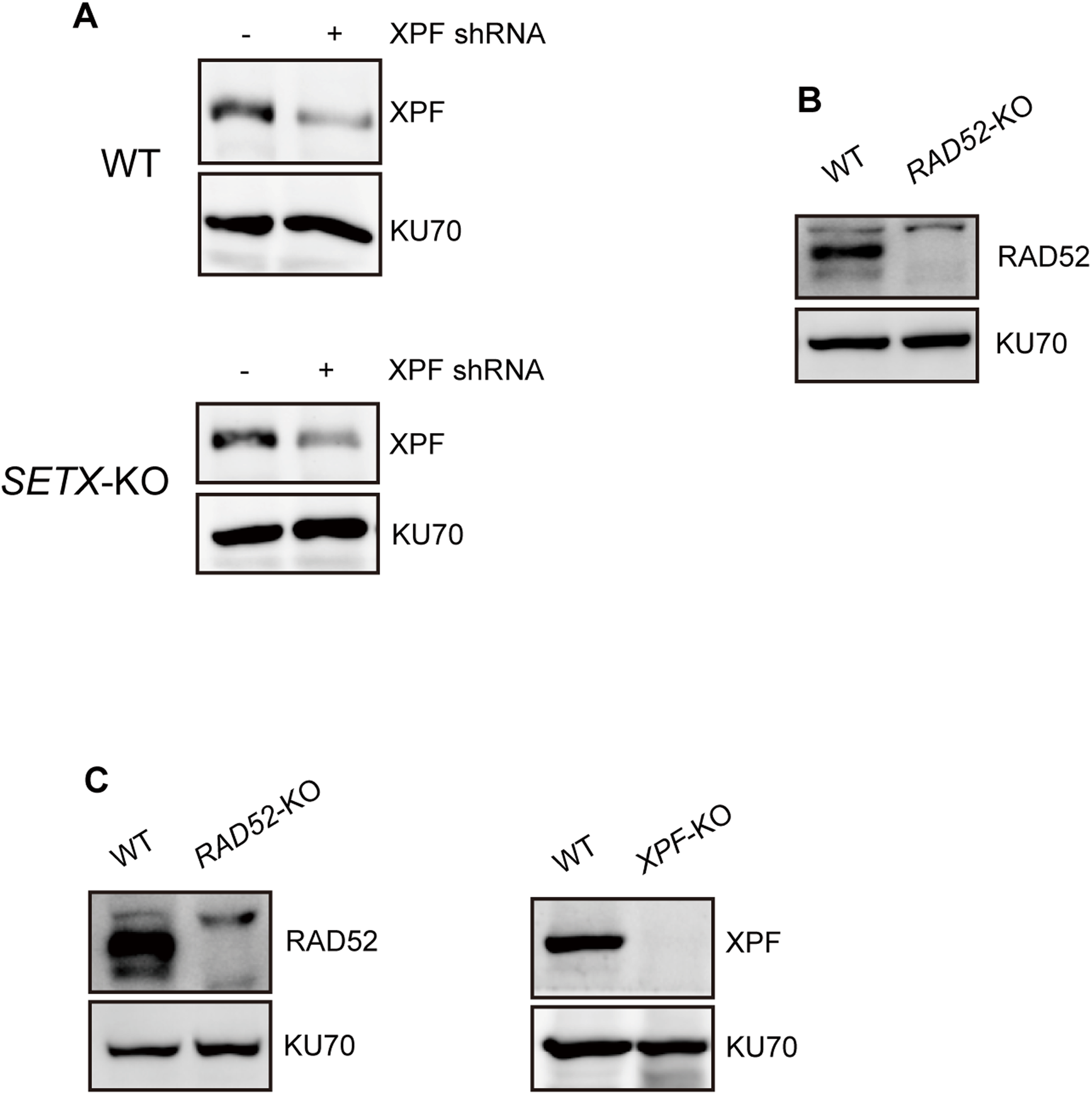
Determining the BIR dependence on XPF/RAD52 in SETX-deficient cells. A. Western blotting showing the efficiency of XPF shRNA in WT and *SETX*-KO U2OS EGFP-BIR-5085 reporter cells. B. Western blotting showing the efficiency of RAD52 knockout in U2OS EGFP-BIR-5085 reporter cells. C. Western blotting showing the efficiency of RAD52 knockout and XPF knockout in U2OS EGFP-BIR-5085 reporter cells.

**Fig S5 related to Fig 5 and Fig 6.**
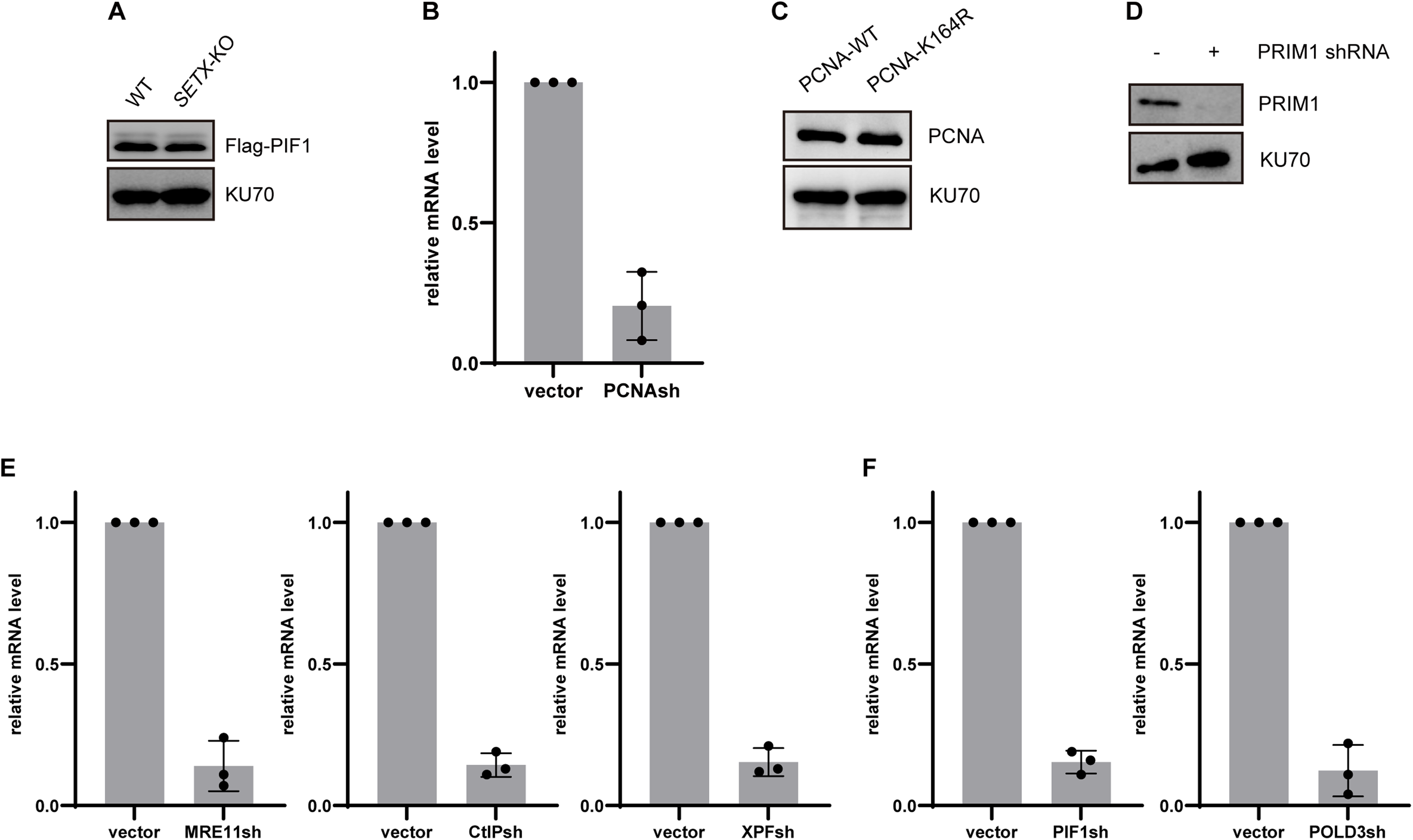
Determining how SETX deficiency activates BIR through PCNA ubiquitination. A. Western blotting showing the expression of Flag-PIF1 in WT and *SETX*-KO U2OS EGFP-BIR-5085 reporter cells. B. qPCR results showing the efficiency of PCNA shRNA in EGFP-BIR-5085 reporter cells. C. Western blotting showing the expression of shRNA-resistant PCNA in WT and *SETX*-KO U2OS EGFP-BIR-5085 reporter cells. D. qPCR results showing the efficiency of PRIM1 shRNA in EGFP-BIR-5085 reporter cells. E. qPCR results showing the efficiency of MRE11 shRNA, CtIP shRNA, and XPF shRNA in U2OS EGFP-HR-5633 reporter cells. F. qPCR results showing the efficiency of PIF1 shRNA and POLD3 shRNA in U2OS EGFP-HR-5633 reporter cells.

**Fig S6.**
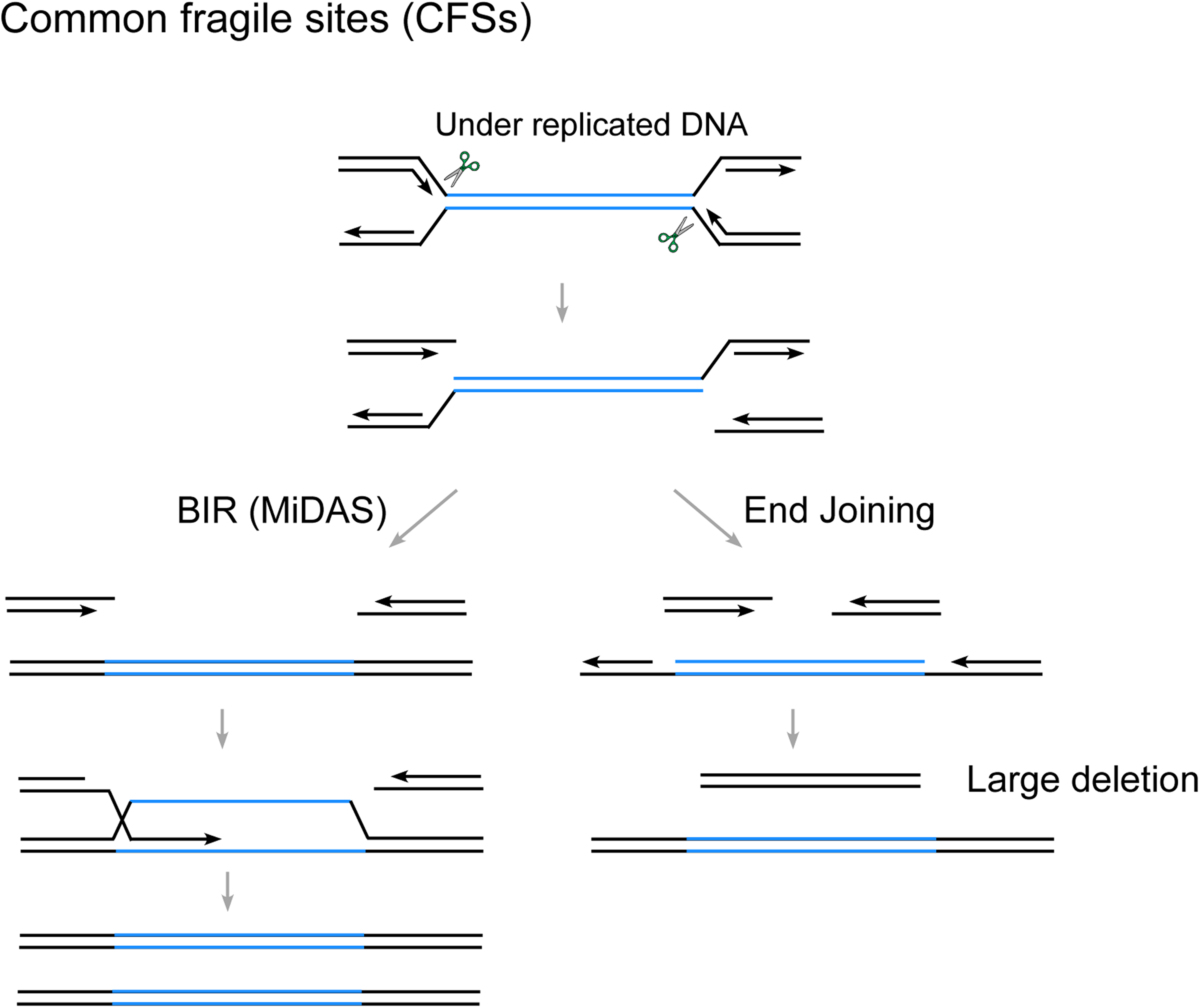
Models showing BIR prevents large deletions at CFSs through MiDAS. Schematic drawing of repair products by BIR (MiDAS) and end joining to repair DSBs at CFSs in mitosis.

## Reference

Alfano, L., Caporaso, A., Altieri, A., Dell’Aquila, M., Landi, C., Bini, L., Pentimalli, F., and Giordano, A. (2019). Depletion of the RNA binding protein HNRNPD impairs homologous recombination by inhibiting DNA-end resection and inducing R-loop accumulation. Nucleic Acids Res 47, 4068–4085.

Alzu, A., Bermejo, R., Begnis, M., Lucca, C., Piccini, D., Carotenuto, W., Saponaro, M., Brambati, A., Cocito, A., Foiani, M., et al. (2012). Senataxin associates with replication forks to protect fork integrity across RNA-polymerase-II-transcribed genes. Cell 151, 835–846.

Anand, R., Jasrotia, A., Bundschuh, D., Howard, S.M., Ranjha, L., Stucki, M., and Cejka, P. (2019). NBS1 promotes the endonuclease activity of the MRE11-RAD50 complex by sensing CtIP phosphorylation. EMBO J 38.

Anand, R., Ranjha, L., Cannavo, E., and Cejka, P. (2016). Phosphorylated CtIP Functions as a Co-factor of the MRE11-RAD50-NBS1 Endonuclease in DNA End Resection. Mol Cell 64, 940–950.

Anand, R.P., Lovett, S.T., and Haber, J.E. (2013). Break-induced DNA replication. Cold Spring Harbor perspectives in biology 5, a010397.

Bennett, C.L., Sopher, B.L., and La Spada, A.R. (2020). Tight expression regulation of senataxin, linked to motor neuron disease and ataxia, is required to avert cell-cycle block and nucleolus disassembly. Heliyon 6, e04165.

Bhowmick, R., Hickson, I.D., and Liu, Y. (2023). Completing genome replication outside of S phase. Mol Cell 83, 3596–3607.

Brickner, J.R., Garzon, J.L., and Cimprich, K.A. (2022). Walking a tightrope: The complex balancing act of R-loops in genome stability. Mol Cell 82, 2267–2297.

Burger, K., Schlackow, M., and Gullerova, M. (2019). Tyrosine kinase c-Abl couples RNA polymerase II transcription to DNA double-strand breaks. Nucleic Acids Res 47, 3467–3484.

Buzovetsky, O., Kwon, Y., Pham, N.T., Kim, C., Ira, G., Sung, P., and Xiong, Y. (2017). Role of the Pif1-PCNA Complex in Pol δ-Dependent Strand Displacement DNA Synthesis and Break-Induced Replication. Cell Reports 21, 1707–1714.

Cannavo, E., and Cejka, P. (2014). Sae2 promotes dsDNA endonuclease activity within Mre11-Rad50-Xrs2 to resect DNA breaks. Nature 514, 122–125.

Ceccaldi, R., Rondinelli, B., and D’Andrea, A.D. (2016). Repair Pathway Choices and Consequences at the Double-Strand Break. Trends Cell Biol 26, 52–64.

Cejka, P., and Symington, L.S. (2021). DNA End Resection: Mechanism and Control. Annu Rev Genet 55, 285–307.

Chang, E.Y., Tsai, S., Aristizabal, M.J., Wells, J.P., Coulombe, Y., Busatto, F.F., Chan, Y.A., Kumar, A., Dan Zhu, Y., Wang, A.Y., et al. (2019). MRE11-RAD50-NBS1 promotes Fanconi Anemia R-loop suppression at transcription-replication conflicts. Nat Commun 10, 4265.

Chen, J.Y., Zhang, X., Fu, X.D., and Chen, L. (2019). R-ChIP for genome-wide mapping of R-loops by using catalytically inactive RNASEH1. Nat Protoc 14, 1661–1685.

Chen, Y.Z., Bennett, C.L., Huynh, H.M., Blair, I.P., Puls, I., Irobi, J., Dierick, I., Abel, A., Kennerson, M.L., Rabin, B.A., et al. (2004). DNA/RNA helicase gene mutations in a form of juvenile amyotrophic lateral sclerosis (ALS4). Am J Hum Genet 74, 1128–1135.

Chen, Y.Z., Hashemi, S.H., Anderson, S.K., Huang, Y., Moreira, M.C., Lynch, D.R., Glass, I.A., Chance, P.F., and Bennett, C.L. (2006). Senataxin, the yeast Sen1p orthologue: characterization of a unique protein in which recessive mutations cause ataxia and dominant mutations cause motor neuron disease. Neurobiol Dis 23, 97–108.

Cohen, S., Guenole, A., Lazar, I., Marnef, A., Clouaire, T., Vernekar, D.V., Puget, N., Rocher, V., Arnould, C., Aguirrebengoa, M., et al. (2022). A POLD3/BLM dependent pathway handles DSBs in transcribed chromatin upon excessive RNA:DNA hybrid accumulation. Nat Commun 13, 2012.

Cohen, S., Puget, N., Lin, Y.L., Clouaire, T., Aguirrebengoa, M., Rocher, V., Pasero, P., Canitrot, Y., and Legube, G. (2018). Senataxin resolves RNA:DNA hybrids forming at DNA double-strand breaks to prevent translocations. Nat Commun 9, 533.

Crossley, M.P., Song, C., Bocek, M.J., Choi, J.H., Kousorous, J., Sathirachinda, A., Lin, C., Brickner, J.R., Bai, G., Lans, H., et al. (2023). R-loop-derived cytoplasmic RNA-DNA hybrids activate an immune response. Nature 613, 187–194.

D’Alessandro, G., Whelan, D.R., Howard, S.M., Vitelli, V., Renaudin, X., Adamowicz, M., Iannelli, F., Jones-Weinert, C.W., Lee, M., Matti, V., et al. (2018). BRCA2 controls DNA:RNA hybrid level at DSBs by mediating RNase H2 recruitment. Nat Commun 9, 5376.

Daley, J.M., Tomimatsu, N., Hooks, G., Wang, W., Miller, A.S., Xue, X., Nguyen, K.A., Kaur, H., Williamson, E., Mukherjee, B., et al. (2020). Specificity of end resection pathways for double-strand break regions containing ribonucleotides and base lesions. Nat Commun 11, 3088.

Deshpande, R.A., Lee, J.H., Arora, S., and Paull, T.T. (2016). Nbs1 Converts the Human Mre11/Rad50 Nuclease Complex into an Endo/Exonuclease Machine Specific for Protein-DNA Adducts. Mol Cell 64, 593–606.

Domingo-Prim, J., Endara-Coll, M., Bonath, F., Jimeno, S., Prados-Carvajal, R., Friedlander, M.R., Huertas, P., and Visa, N. (2019). EXOSC10 is required for RPA assembly and controlled DNA end resection at DNA double-strand breaks. Nat Commun 10, 2135.

Donnianni, R.A., and Symington, L.S. (2013). Break-induced replication occurs by conservative DNA synthesis. Proc Natl Acad Sci U S A 110, 13475–13480.

Gao, M., Wei, W., Li, M.M., Wu, Y.S., Ba, Z., Jin, K.X., Li, M.M., Liao, Y.Q., Adhikari, S., Chong, Z., et al. (2014). Ago2 facilitates Rad51 recruitment and DNA double-strand break repair by homologous recombination. Cell Res 24, 532–541.

Garcia-Muse, T., and Aguilera, A. (2019). R Loops: From Physiological to Pathological Roles. Cell 179, 604–618.

Gatti, V., De Domenico, S., Melino, G., and Peschiaroli, A. (2023). Senataxin and R-loops homeostasis: multifaced implications in carcinogenesis. Cell Death Discov 9, 145.

Giannini, M., and Porrua, O. (2024). Senataxin: A key actor in RNA metabolism, genome integrity and neurodegeneration. Biochimie 217, 10–19.

Gomez-Gonzalez, B., and Aguilera, A. (2023). Break-induced RNA-DNA hybrids (BIRDHs) in homologous recombination: friend or foe? EMBO Rep 24, e57801.

Groh, M., Albulescu, L.O., Cristini, A., and Gromak, N. (2017). Senataxin: Genome Guardian at the Interface of Transcription and Neurodegeneration. J Mol Biol 429, 3181–3195.

Han, T., Goralski, M., Capota, E., Padrick, S.B., Kim, J., Xie, Y., and Nijhawan, D. (2016). The antitumor toxin CD437 is a direct inhibitor of DNA polymerase alpha. Nat Chem Biol 12, 511–515.

Hatchi, E., Skourti-Stathaki, K., Ventz, S., Pinello, L., Yen, A., Kamieniarz-Gdula, K., Dimitrov, S., Pathania, S., McKinney, K.M., Eaton, M.L., et al. (2015). BRCA1 recruitment to transcriptional pause sites is required for R-loop-driven DNA damage repair. Mol Cell 57, 636–647.

Helmrich, A., Ballarino, M., and Tora, L. (2011). Collisions between replication and transcription complexes cause common fragile site instability at the longest human genes. Mol Cell 44, 966–977.

Heyer, W.D. (2015). Regulation of recombination and genomic maintenance. Cold Spring Harbor perspectives in biology 7, a016501.

Hustedt, N., and Durocher, D. (2016). The control of DNA repair by the cell cycle. Nat Cell Biol 19, 1–9.

In, S., Renck Nunes, P., Valador Fernandes, R., and Lingner, J. (2025). TERRA R-loops trigger a switch in telomere maintenance towards break-induced replication and PrimPol-dependent repair. bioRxiv.

Jain, S., Sugawara, N., Lydeard, J., Vaze, M., Tanguy Le Gac, N., and Haber, J.E. (2009). A recombination execution checkpoint regulates the choice of homologous recombination pathway during DNA double-strand break repair. Genes Dev 23, 291–303.

Jang, Y., Elsayed, Z., Eki, R., He, S., Du, K.P., Abbas, T., and Kai, M. (2020). Intrinsically disordered protein RBM14 plays a role in generation of RNA:DNA hybrids at double-strand break sites. Proc Natl Acad Sci U S A 117, 5329–5338.

Jasin, M., and Rothstein, R. (2013). Repair of strand breaks by homologous recombination. Cold Spring Harbor perspectives in biology 5.

Khanna, K.K., and Jackson, S.P. (2001). DNA double-strand breaks-signaling, repair and the cancer connection. Nature Genet 27, 247–254.

Kramara, J., Osia, B., and Malkova, A. (2018). Break-Induced Replication: The Where, The Why, and The How. Trends Genet 34, 518–531.

Li, L., Germain, D.R., Poon, H.Y., Hildebrandt, M.R., Monckton, E.A., McDonald, D., Hendzel, M.J., and Godbout, R. (2016). DEAD Box 1 Facilitates Removal of RNA and Homologous Recombination at DNA Double-Strand Breaks. Mol Cell Biol 36, 2794–2810.

Li, S., Wang, H., Jehi, S., Li, J., Liu, S., Wang, Z., Truong, L., Chiba, T., Wang, Z., and Wu, X. (2021a). PIF1 helicase promotes break-induced replication in mammalian cells. EMBO J, e104509.

Li, S., Wang, H., Jehi, S., Li, J., Liu, S., Wang, Z., Truong, L., Chiba, T., Wang, Z., and Wu, X. (2021b). PIF1 helicase promotes break-induced replication in mammalian cells. EMBO J 40, e104509.

Liaw, H., Lee, D., and Myung, K. (2011). DNA-PK-dependent RPA2 hyperphosphorylation facilitates DNA repair and suppresses sister chromatid exchange. PLoS One 6, e21424.

Liu, L., and Malkova, A. (2022). Break-induced replication: unraveling each step. Trends Genet 38, 752–765.

Liu, S., Hua, Y., Wang, J., Li, L., Yuan, J., Zhang, B., Wang, Z., Ji, J., and Kong, D. (2021). RNA polymerase III is required for the repair of DNA double-strand breaks by homologous recombination. Cell 184, 1314–1329 e1310.

Liu, S., Wang, Z., Shah, S.B., Chang, C.Y., Ai, M., Nguyen, T., Xiang, R., and Wu, X. (2023). DNA repair protein RAD52 is required for protecting G-quadruplexes in mammalian cells. J Biol Chem 299, 102770.

Llorente, B., Smith, C.E., and Symington, L.S. (2008). Break-induced replication: what is it and what is it for? Cell Cycle 7, 859–864.

Lydeard, J.R., Jain, S., Yamaguchi, M., and Haber, J.E. (2007). Break-induced replication and telomerase-independent telomere maintenance require Pol32. Nature 448, 820–823.

Makharashvili, N., Arora, S., Yin, Y., Fu, Q., Wen, X., Lee, J.H., Kao, C.H., Leung, J.W., Miller, K.M., and Paull, T.T. (2018). Sae2/CtIP prevents R-loop accumulation in eukaryotic cells. Elife 7.

Marnef, A., and Legube, G. (2021). R-loops as Janus-faced modulators of DNA repair. Nat Cell Biol 23, 305–313.

Matsui, M., Sakasai, R., Abe, M., Kimura, Y., Kajita, S., Torii, W., Katsuki, Y., Ishiai, M., Iwabuchi, K., Takata, M., et al. (2020). USP42 enhances homologous recombination repair by promoting R-loop resolution with a DNA-RNA helicase DHX9. Oncogenesis 9, 60.

Mehta, A., Beach, A., and Haber, J.E. (2017). Homology Requirements and Competition between Gene Conversion and Break-Induced Replication during Double-Strand Break Repair. Mol Cell 65, 515–526 e513.

Michelini, F., Pitchiaya, S., Vitelli, V., Sharma, S., Gioia, U., Pessina, F., Cabrini, M., Wang, Y., Capozzo, I., Iannelli, F., et al. (2017). Damage-induced lncRNAs control the DNA damage response through interaction with DDRNAs at individual double-strand breaks. Nat Cell Biol 19, 1400–1411.

Mirman, Z., Lottersberger, F., Takai, H., Kibe, T., Gong, Y., Takai, K., Bianchi, A., Zimmermann, M., Durocher, D., and de Lange, T. (2018). 53BP1-RIF1-shieldin counteracts DSB resection through CST- and Polalpha-dependent fill-in. Nature 560, 112–116.

Mirman, Z., Sasi, N.K., King, A., Chapman, J.R., and de Lange, T. (2022). 53BP1-shieldin-dependent DSB processing in BRCA1-deficient cells requires CST-Polalpha-primase fill-in synthesis. Nat Cell Biol 24, 51–61.

Moreira, M.C., Klur, S., Watanabe, M., Nemeth, A.H., Le Ber, I., Moniz, J.C., Tranchant, C., Aubourg, P., Tazir, M., Schols, L., et al. (2004). Senataxin, the ortholog of a yeast RNA helicase, is mutant in ataxia-ocular apraxia 2. Nat Genet 36, 225–227.

Motycka, T.A., Bessho, T., Post, S.M., Sung, P., and Tomkinson, A.E. (2004). Physical and functional interaction between the XPF/ERCC1 endonuclease and hRad52. J Biol Chem 279, 13634–13639.

Niehrs, C., and Luke, B. (2020). Regulatory R-loops as facilitators of gene expression and genome stability. Nat Rev Mol Cell Biol 21, 167–178.

Niimi, A., Brown, S., Sabbioneda, S., Kannouche, P.L., Scott, A., Yasui, A., Green, C.M., and Lehmann, A.R. (2008). Regulation of proliferating cell nuclear antigen ubiquitination in mammalian cells. Proc Natl Acad Sci U S A 105, 16125–16130.

Ohle, C., Tesorero, R., Schermann, G., Dobrev, N., Sinning, I., and Fischer, T. (2016). Transient RNA-DNA Hybrids Are Required for Efficient Double-Strand Break Repair. Cell 167, 1001–1013 e1007.

Paques, F., and Haber, J.E. (1999). Multiple pathways of recombination induced by double-strand breaks in Saccharomyces cerevisiae. Microbiol Mol Biol Rev 63, 349–404.

Petermann, E., Lan, L., and Zou, L. (2022). Sources, resolution and physiological relevance of R-loops and RNA-DNA hybrids. Nat Rev Mol Cell Biol 23, 521–540.

Pham, N., Yan, Z., Yu, Y., Faria Afreen, M., Malkova, A., Haber, J.E., and Ira, G. (2021). Mechanisms restraining break-induced replication at two-ended DNA double-strand breaks. EMBO J 40, e104847.

Rawal, C.C., Zardoni, L., Di Terlizzi, M., Galati, E., Brambati, A., Lazzaro, F., Liberi, G., and Pellicioli, A. (2020). Senataxin Ortholog Sen1 Limits DNA:RNA Hybrid Accumulation at DNA Double-Strand Breaks to Control End Resection and Repair Fidelity. Cell Rep 31, 107603.

Rhodes, D.R., Kalyana-Sundaram, S., Mahavisno, V., Varambally, R., Yu, J., Briggs, B.B., Barrette, T.R., Anstet, M.J., Kincead-Beal, C., Kulkarni, P., et al. (2007). Oncomine 3.0: genes, pathways, and networks in a collection of 18,000 cancer gene expression profiles. Neoplasia 9, 166–180.

Richard, P., Feng, S., and Manley, J.L. (2013). A SUMO-dependent interaction between Senataxin and the exosome, disrupted in the neurodegenerative disease AOA2, targets the exosome to sites of transcription-induced DNA damage. Genes Dev 27, 2227–2232.

Said, M., Barra, V., Balzano, E., Talhaoui, I., Pelliccia, F., Giunta, S., and Naim, V. (2022). FANCD2 promotes mitotic rescue from transcription-mediated replication stress in SETX-deficient cancer cells. Commun Biol 5, 1395.

Saini, N., Ramakrishnan, S., Elango, R., Ayyar, S., Zhang, Y., Deem, A., Ira, G., Haber, J.E., Lobachev, K.S., and Malkova, A. (2013). Migrating bubble during break-induced replication drives conservative DNA synthesis. Nature 502, 389–392.

Sakofsky, C.J., Ayyar, S., Deem, A.K., Chung, W.H., Ira, G., and Malkova, A. (2015). Translesion Polymerases Drive Microhomology-Mediated Break-Induced Replication Leading to Complex Chromosomal Rearrangements. Mol Cell 60, 860–872.

Sakofsky, C.J., Ayyar, S., and Malkova, A. (2012). Break-induced replication and genome stability. Biomolecules 2, 483–504.

Sakofsky, C.J., Roberts, S.A., Malc, E., Mieczkowski, P.A., Resnick, M.A., Gordenin, D.A., and Malkova, A. (2014). Break-induced replication is a source of mutation clusters underlying kataegis. Cell Rep 7, 1640–1648.

Sessa, G., Gomez-Gonzalez, B., Silva, S., Perez-Calero, C., Beaurepere, R., Barroso, S., Martineau, S., Martin, C., Ehlen, A., Martinez, J.S., et al. (2021). BRCA2 promotes DNA-RNA hybrid resolution by DDX5 helicase at DNA breaks to facilitate their repairdouble dagger. EMBO J 40, e106018.

Shah, S.B., Li, Y., Li, S., Hu, Q., Wu, T., Shi, Y., Nguyen, T., Ive, I., Shi, L., Wang, H., et al. (2024). 53BP1 deficiency leads to hyperrecombination using break-induced replication (BIR). Nat Commun 15, 8648.

Shibata, A., Moiani, D., Arvai, A.S., Perry, J., Harding, S.M., Genois, M.M., Maity, R., van Rossum-Fikkert, S., Kertokalio, A., Romoli, F., et al. (2014). DNA double-strand break repair pathway choice is directed by distinct MRE11 nuclease activities. Mol Cell 53, 7–18.

Smith, C.E., Llorente, B., and Symington, L.S. (2007). Template switching during break-induced replication. Nature 447, 102–105.

Sollier, J., Stork, C.T., Garcia-Rubio, M.L., Paulsen, R.D., Aguilera, A., and Cimprich, K.A. (2014). Transcription-coupled nucleotide excision repair factors promote R-loop-induced genome instability. Mol Cell 56, 777–785.

Stork, C.T., Bocek, M., Crossley, M.P., Sollier, J., Sanz, L.A., Chedin, F., Swigut, T., and Cimprich, K.A. (2016). Co-transcriptional R-loops are the main cause of estrogen-induced DNA damage. Elife 5.

Suraweera, A., Lim, Y., Woods, R., Birrell, G.W., Nasim, T., Becherel, O.J., and Lavin, M.F. (2009). Functional role for senataxin, defective in ataxia oculomotor apraxia type 2, in transcriptional regulation. Human molecular genetics 18, 3384–3396.

Tan, J., Duan, M., Yadav, T., Phoon, L., Wang, X., Zhang, J.M., Zou, L., and Lan, L. (2020). An R-loop-initiated CSB-RAD52-POLD3 pathway suppresses ROS-induced telomeric DNA breaks. Nucleic Acids Res 48, 1285–1300.

Thakar, T., Leung, W., Nicolae, C.M., Clements, K.E., Shen, B., Bielinsky, A.K., and Moldovan, G.L. (2020). Ubiquitinated-PCNA protects replication forks from DNA2-mediated degradation by regulating Okazaki fragment maturation and chromatin assembly. Nat Commun 11, 2147.

Ursic, D., Chinchilla, K., Finkel, J.S., and Culbertson, M.R. (2004). Multiple protein/protein and protein/RNA interactions suggest roles for yeast DNA/RNA helicase Sen1p in transcription, transcription-coupled DNA repair and RNA processing. Nucleic Acids Res 32, 2441–2452.

Wilson, M.A., Kwon, Y., Xu, Y., Chung, W.H., Chi, P., Niu, H., Mayle, R., Chen, X., Malkova, A., Sung, P., et al. (2013). Pif1 helicase and Poldelta promote recombination-coupled DNA synthesis via bubble migration. Nature 502, 393–396.

Wu, X., and Malkova, A. (2021a). Break-induced replication mechanisms in yeast and mammals. Curr Opin Genet Dev 71, 163–170.

Wu, X., and Malkova, A. (2021b). Break-induced replication mechanisms in yeast and mammals. Curr Opin Genet Dev 71, 163–170.

Yadav, T., Zhang, J.M., Ouyang, J., Leung, W., Simoneau, A., and Zou, L. (2022). TERRA and RAD51AP1 promote alternative lengthening of telomeres through an R- to D-loop switch. Mol Cell 82, 3985–4000 e3984.

Yasuhara, T., Kato, R., Hagiwara, Y., Shiotani, B., Yamauchi, M., Nakada, S., Shibata, A., and Miyagawa, K. (2018). Human Rad52 Promotes XPG-Mediated R-loop Processing to Initiate Transcription-Associated Homologous Recombination Repair. Cell 175, 558–570 e511.

Yu, Z., Mersaoui, S.Y., Guitton-Sert, L., Coulombe, Y., Song, J., Masson, J.Y., and Richard, S. (2020). DDX5 resolves R-loops at DNA double-strand breaks to promote DNA repair and avoid chromosomal deletions. NAR Cancer 2, zcaa028.

Yuce, O., and West, S.C. (2013). Senataxin, defective in the neurodegenerative disorder ataxia with oculomotor apraxia 2, lies at the interface of transcription and the DNA damage response. Mol Cell Biol 33, 406–417.

Zdravkovic, A., Daley, J.M., Dutta, A., Niwa, T., Murayama, Y., Kanamaru, S., Ito, K., Maki, T., Argunhan, B., Takahashi, M., et al. (2021). A conserved Ctp1/CtIP C-terminal peptide stimulates Mre11 endonuclease activity. Proc Natl Acad Sci U S A 118.

Zhao, B., Rothenberg, E., Ramsden, D.A., and Lieber, M.R. (2020). The molecular basis and disease relevance of non-homologous DNA end joining. Nat Rev Mol Cell Biol 21, 765–781.

Zhao, H., Hartono, S.R., de Vera, K.M.F., Yu, Z., Satchi, K., Zhao, T., Sciammas, R., Sanz, L., Chedin, F., and Barlow, J. (2022). Senataxin and RNase H2 act redundantly to suppress genome instability during class switch recombination. Elife 11.

Zhou, Y., Caron, P., Legube, G., and Paull, T.T. (2014). Quantitation of DNA double-strand break resection intermediates in human cells. Nucleic Acids Res 42, e19.

